# TAK1 blockade as a therapy for retinal neovascularization

**DOI:** 10.1101/2021.01.29.428701

**Authors:** Fan-Li Lin, Jiang-Hui Wang, Jinying Chen, Linxin Zhu, Yu-Fan Chuang, Leilei Tu, Chenkai Ma, Suraj Lama, Damien Ling, Raymond Ching-Bong Wong, Alex W. Hewitt, Ching-Li Tseng, Bang V. Bui, Peter van Wijngaarden, Gregory J. Dusting, Peng-Yuan Wang, Guei-Sheung Liu

## Abstract

Retinal neovascularization, or pathological angiogenesis in the retina, is a leading cause of blindness in developed countries. Transforming growth factor-β-activated kinase 1 (TAK1) is a mitogen-activated protein kinase kinase kinase (MAPKKK) activated by TGF-β1 and other pro-inflammatory cytokines. TAK1 is also a key mediator of inflammation, innate immune responses, apoptosis and tissue homeostasis and plays an important role in physiological angiogenesis. Its role in pathological angiogenesis, particularly in retinal neovascularization, remains unclear. We investigated the regulatory role of TAK1 in pathological angiogenesis in the retina. Transcriptome analysis of human retina featuring retinal neovascularization revealed enrichment of known TAK1-mediated signaling pathways. Selective inhibition of TAK1 activation by 5Z-7-oxozeaenol attenuated aberrant retinal angiogenesis in rats following oxygen-induced retinopathy. Transcriptome profiling revealed that TAK1 activation in human microvascular endothelial cells under TNFα stimulation led to increase the gene expression related to cytokines and leukocyte-endothelial interaction, mainly through nuclear factor kappa B (NFκB) signaling pathways. These results reveal that inhibition of TAK1 signaling may have therapeutic value for the treatment of pathological angiogenesis in the retina.

## INTRODUCTION

Retinal neovascularization is a severe complication of several ocular diseases, often resulting in permanent damage to the neurosensory retina and irreversible vision loss (1). Retinal neovascularization shares similar characteristics to pathological angiogenesis elsewhere in the body, exhibiting features including endothelial proliferation and migration, elevated vascular permeability and inflammation; all processes in which vascular endothelial growth factor (VEGF) plays a crucial role (2). Recently, the advent of ocular anti-VEGF therapies has been a significant advance in mitigating the risk of vison loss associated with retinal neovascularization. Despite this success, there is increasing evidence that some patients do not respond well to anti-VEGF treatments (3). Therefore, it is necessary to seek other therapeutic targets.

The common pathways shared between inflammation and angiogenesis play critical roles in retinal neovascularization (4). Several studies have reported increased expression of various pro-inflammatory cytokines, chemokines, pro-angiogenic factors and adhesion molecules in patients with retinal neovascularization, such as tumor necrosis factor α (TNFα), interleukin-1 (IL-1), chemokine (C-X-C motif) ligand 8 (CXCL8; known as IL-8), vascular endothelial growth factor (VEGF), as well as intercellular adhesion molecule 1 (ICAM-1 and VCAM-1) (5–9). TNFα was found to suppress the expression of tight junction proteins and is required for VEGF-mediated hyperpermeability in endothelial cells and the breakdown of the blood-retinal barrier in patients with diabetic retinopathy (DR) (10, 11). Interleukins are also primary mediators of inflammation that can promote angiogenesis via induction of proangiogenic factors such as VEGF (4). Moreover, TNFα and IL-1 stimulates expression of ICAM-1, CXCL8 and VEGF in endothelial cells and microglia cells, both of which promote progression of retinal neovascularization (12, 13). A key role for inflammatory mediators is highlighted by their growing use as prognostic markers of the severity of proliferative diabetic retinopathy (PDR) (4).

Transforming growth factor-β-activated kinase 1 (TAK1), a member of the mitogen-activated protein kinase (MAPK) family, is a critical serine/threonine kinase in several cellular signaling pathways. Such pathways can be activated by diverse pro-inflammatory stimuli including TNFα, IL-1, transforming growth factor beta (TGF-β), or toll-like receptor (TLR) ligands. The engagement of TAK1 in turn activates downstream signaling pathways, including nuclear factor kappa B (NFκB) and MAPK p38, extracellular signal-regulated kinase (ERK), and c-Jun kinase (JNK) signaling, modulating inflammatory responses and cell survival (14–16). TAK1 inhibition has been shown to reduce TNFα and CXCL8 secretion in leukocytes (17) and to suppress prostaglandin-endoperoxide synthase 2 (PTGS2; known as cyclooxygenase-2, COX-2), ICAM-1, and VEGF expression in cancer cells through NFκB signaling pathway (18). Expression of TAK1 provides protection for endothelial cells against TNFα-induced apoptosis under inflammatory conditions (19). Indeed, TAK1 deletion causes embryonic lethality as the capacity to prevent TNFα-induced endothelial cell death and vessel regression is lost. Moreover, TAK1 is also important for supporting TNFα-independent vascular formation and endothelial migration. These broad roles for TAK1 suggests that endothelial TAK1 inhibition may be a useful alternate anti-angiogenesis target in retinal disease (19, 20).

We hypothesized that TAK1 plays a crucial role in pathological angiogenesis and TAK1 inhibition can suppress retinal neovascularization. Our findings indicate that TAK1 is an important regulator of inflammatory and angiogenesis-related pathways that contribute to retinal neovascularization. Pharmacological inhibition of TAK1 alleviated aberrant retinal angiogenesis by modulating cytokine signaling and leukocyte-endothelium interactions in a rat model of ischemia-induced retinopathy.

## RESULTS

### TAK1 is expressed in human retina and is positively associated with retinal neovascularization

To understand TAK1 expression profiles in the human retina, we first analyzed published single nuclei RNAseq (snRNAseq, GSE135133) data to examine TAK1 expression in different retinal cells. Analysis of 121,799 individual cells from human retina revealed that both *TAK1* (*MAP3K7*) and its binding protein, *TAB1-3,* are expressed moderately in most neural and non-neural retinal cells (**Fig. 1A and S1**). Notably, among the retinal cells examined, TAK1 is highly enriched in endothelial cells. Immunostaining suggests that TAK1 is expressed in various layers of the human retina in both neural and non-neural cells, such as mural cells and smooth muscle cells (**Fig. 1B**) in larger vessels (diameter of artery or vein ranging from 0.6 - 16 mm) (21). We also found that co-localization of TAK1 and endothelial cell marker CD31 was more likely to be found in the microvasculature, including arterioles or venules (20 - 25 μm) and capillaries (9 μm).

**Fig. 1.**
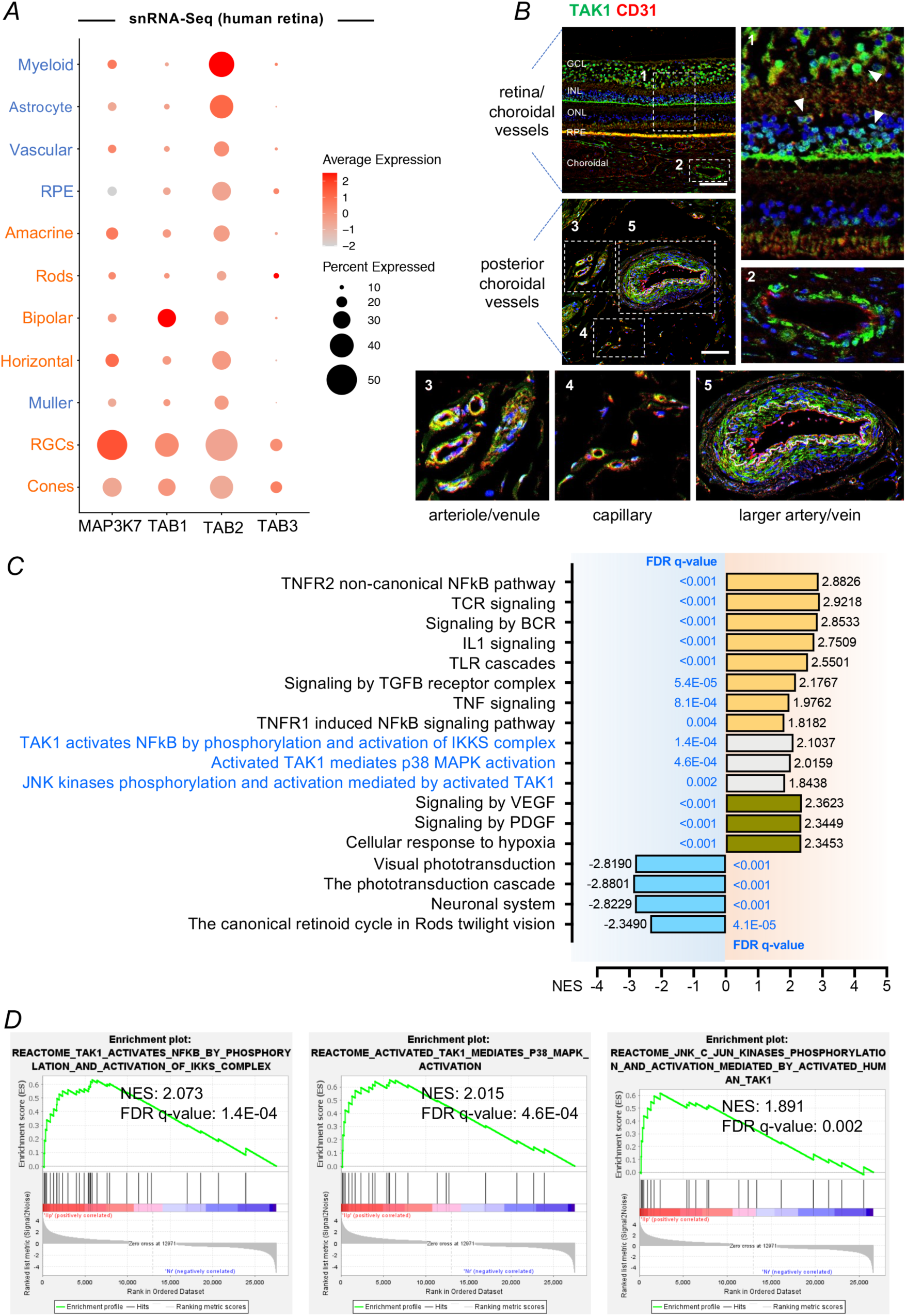
Characterization of TAK1 expression in patients with retinal neovascularization associated with proliferative diabetic retinopathy. (A) Single nuclei RNAseq (snRNAseq) characterization of TAK1 (MAP3K7) expression in normal human retina. TAK1 and TAK1-associated proteins (TAB1-3) were expressed in most neural (red) and non-neural (blue) cells in the retina, including cell associated with the retinal vasculature. (B) Characterization of TAK1 expression in the normal human retina. Endothelial cells were labelled with CD31 (red) and counterstained with TAK1 (green) in retinal section. Apart from neural cells, TAK1 expression was also found in vascular endothelial cells, especially in retinal capillaries (3 and 4). Scale bar: 100 µm. (C) Gene-set enrichment analysis (GSEA) indicated that TAK1 upstream pathways including IL1, TNF, TCR, BCR, TLR, TNFR, or TGFβ signaling (Reactome Pathway Database) were positively enriched in retinal samples from patients with retinal neovascularization due to proliferative diabetic retinopathy. Engagement of TAK1-activated NFκB, p38 MAPK and JNK pathways as well as negative impact on retinal neural functions were also identified. (D) GSEA Enrichment plots illustration of gene set involved in TAK1-NFκB, TAK1-p38 MAPK and TAK1-JNK pathways in the retina of patients with proliferative diabetic retinopathy. GCL: ganglion cell layer; INL: inner nuclear layer; ONL: outer nuclear layer; RPE: retinal pigment epithelium; NSE: normalized enrichment score; FDR: false discovery rate.

To identify the key molecular signaling or biological pathways relevant to the angiogenic process in patients with retinal neovascularization, a transcriptome dataset from humans with proliferative diabetic retinopathy (PDR) was interrogated by taking advantage of Gene Set Enrichment Analysis (GSEA), a knowledge-based approach for interpreting genome-wide expression profiles (22). Compared with controls, those with retinal neovascularization showed enrichment of known TAK1 upstream signaling pathways, including IL1, TNF, TGF-β, TLR, T-cell receptor (TCR) and B-cell receptor (BCR) (**Fig. 1C and S2**). Pathways related directly to TAK1 were enriched, such as “TAK1 activation of NF*κ*B by phosphorylation and activation of inhibitor of κB (IκB) kinase (IKK) complex”, “activated TAK1 mediated p38 MAPK activation”, and “JNK kinases phosphorylation and activation mediated by activated TAK1” (**Fig. 1C and 1D**). In general, genes in association with TAK1 activation mediated by NF*κ*B and MAPKs phosphorylation were up-regulated in patients with retinal neovascularization compared with controls (**Fig. S3**). There was negative enrichment of genes associated with several biological pathways in relation to visual phototransduction and neuronal systems (**Fig 1C**).

### TAK1 is up-regulated in neovascular tufts and positively associated with retinal neovascularization in ischemia-induced retinopathy in rat

To investigate the role of TAK1 in pathological neovascularization in the retina, we subsequently used a rat model of oxygen-induced retinopathy (OIR) which yields ischemic avascular zones and preretinal neovascularization similar to that observed clinically in PDR and retinopathy of prematurity. Neonatal rats were subjected to daily cycles of 80% oxygen for 21 hours and ambient air for 3 hours from postnatal day 0 (P0) to P14 to induce vaso-obliteration. At P14 rats were returned to room air, where maximal preretinal neovascularization occurred by P18, which was followed by a phase of vascular regression (**Fig. 2A**). A significant increase in TAK1 expression at both RNA and protein levels was observed in OIR retinae at P18 compared with controls (**Fig. 2B and 2C**). Examination of the retinal distribution of TAK1 expression revealed that TAK1 expression was uniquely expressed in areas of active neovascularization (**Fig. 2D**). To gain further insight into the role of TAK1 activation at the peak of retinal neovascularization (P18), we then performed transcriptome analyses using bulk-RNA sequencing. GSEA-based analysis revealed a positive enrichment of several TAK1 upstream pathways, including IL1, TNF, TCR, BCR and TLR (**Fig. 2E and S4**), which is consistent with our findings in patients with retinal neovascularization due to PDR. Similar to findings in humans, there was a negative enrichment of genes associated with phototransduction and neuronal systems at P18 in the rats. Of note, certain canonical pathways and biological processes related to angiogenesis were positively enriched in the OIR model, such as VEGF-A/VEGFR2, PDGF pathway, the cellular response to hypoxia, angiogenesis and inflammation (**Fig. 2E and S4**). Overall, our data implicates active involvement of TAK1 in retinal neovascularization.

**Fig. 2.**
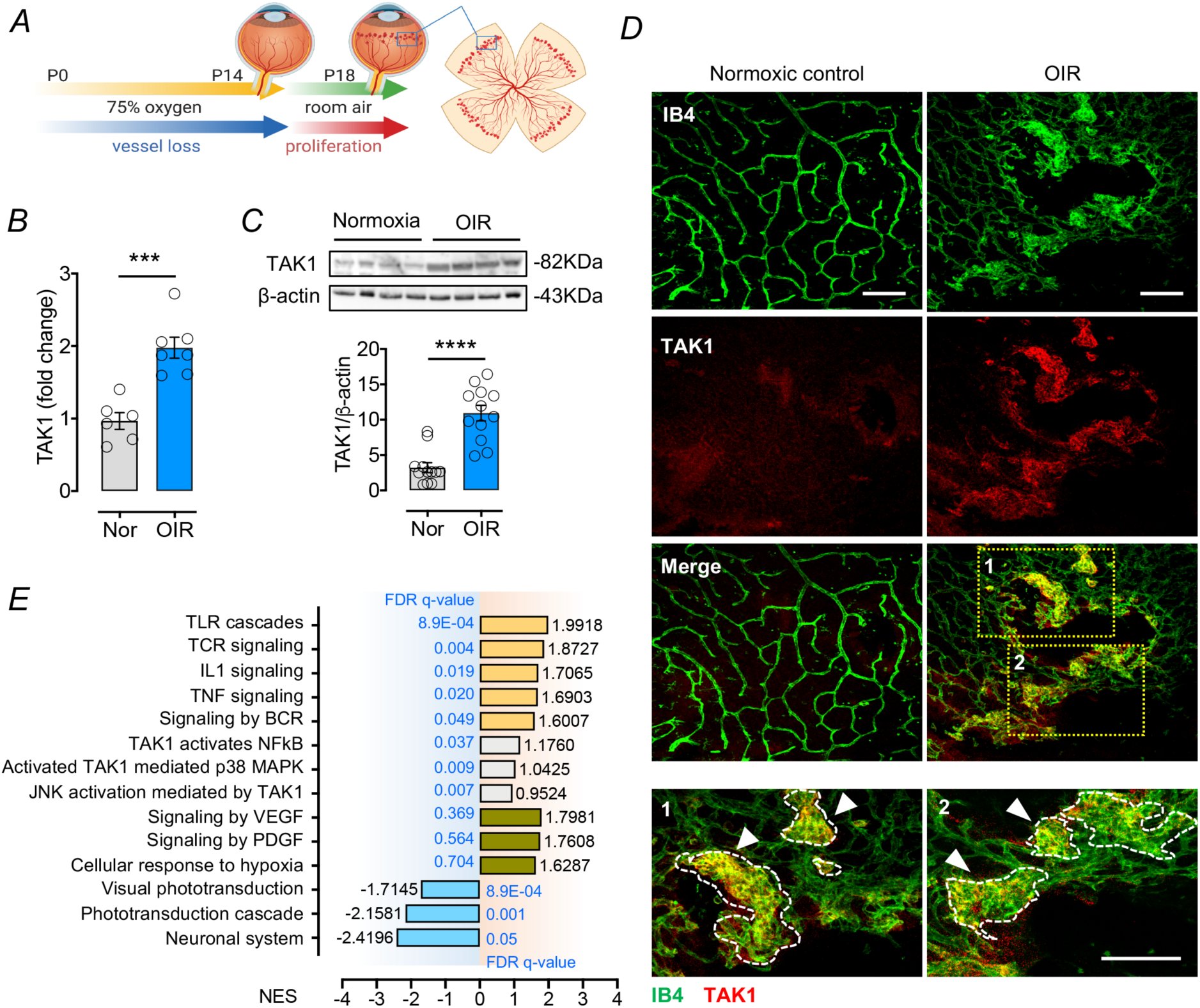
Involvement of TAK1 in pathological retinal angiogenesis in a rat model of oxygen-induced retinopathy. (A) Schematic diagram of the Sprague-Dawley rat model of oxygen-induced retinopathy (OIR). Neonatal rats were exposed to 80% of oxygen for 21 hours each day from postnatal day 0 (P0) to P14. Rats were returned to room air from P14 to P18. Vessel loss was induced during oxygen exposure and retinal angiogenesis could be induced after rats returned to room air. (B) qPCR analysis of *TAK1* expression in OIR at P18 as compared with the normoxic control (Nor) group (n = 6-7). (C) Western blot analysis of TAK1 expression in OIR at P18 compared with normoxic control group (n = 10-12). Representative immunoblotting from 4 independent samples were shown for normoxic control (lanes 1-4) and OIR group (lanes 5-8). (D) Immunobiological staining indicated expression of TAK1 in the pathological vessel. Vasculature were visualized by isolectin GS-IB4 labelling (green) and counterstained with TAK1 (red) in retinal whole mount. Arrowheads indicate vessel tufts. Scale bar: 100 µm. (E) GSEA indicated that most of TAK1 upstream pathways including TNF, IL1, TCR, BCR, TLR, and VEGF pathways (from Reactome Pathway Database) were positively enriched in OIR at P18. Engagement of TAK1-activated NFκB, p38 MAPK and JNK pathways as well as negative impact on retinal neural functions were also identified. Group data are shown as means ± SEM. Statistical analysis was undertaken with two-tailed unpaired t-test; \*\*\**P* < 0.001, \*\*\*\**P* < 0.0001. NSE: normalized enrichment score; FDR: false discovery rate.

### Inhibition of TAK1 activation attenuates retinal neovascularization in rat OIR model

Given that the ablation of TAK1 in endothelial cells impairs angiogenesis and causes lethality (19, 20), we therefore used a selective inhibitor of TAK1, 5Z-7-oxozeaenol, a resorcylic acid lactone derived from a fungus (23), to assess whether TAK1 signaling blockade can alleviate retinal neovascularization. 5Z-7-oxozeaenol was administered intravitreally to the OIR rats at P14 (**Fig. 3A**). Four days after drug administration, retinal neovascularization was significantly suppressed by 51.3% and 57.1% in OIR rats that had received low (18 ng) and high doses (90 ng) of 5Z-7-oxozeaenol, respectively, compared to OIR rats that had vehicle only (**Fig. 3B and S5**). No significant difference in retinal vaso-obliteration was found among these groups (**Fig. 3C**). We also examined the development of deep capillary beds to assess the effects of TAK1 inhibition in physiological vessel growth (24). Normal control rats developed a complete deep layer vasculature, while such development was severely disrupted in OIR rats (**Fig. 3D**). We found that 5Z-7-oxozeaenol treatment did not improve or damage the formation of deep capillary beds in OIR retina, which is likely attributed to suppression of proliferative vessels by TAK1 inhibition. Moreover, the rats receiving low or high dose of 5Z-7-oxozeaenol did not show any abnormalities in the retinal vasculature under normoxic conditions (**Fig. S6**).

**Fig. 3.**
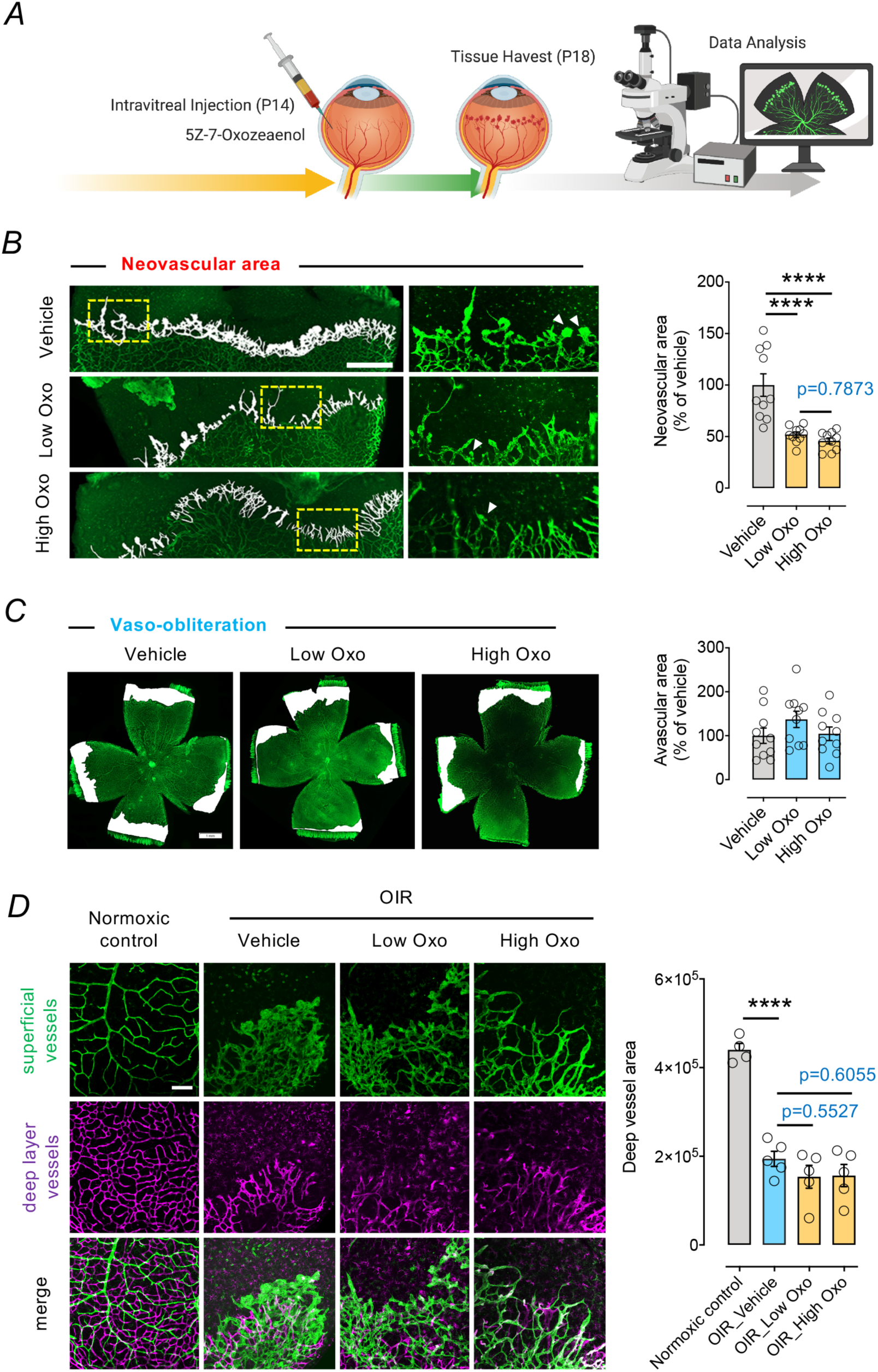
TAK1 Inhibition by 5Z-7-oxozeaenol suppressed retinal angiogenesis in OIR rats. (A) Experimental protocol for 5Z-7-oxozeaenol treatment in the OIR rat model. A single intravitreal injection of vehicle, low (18 ng) or high (90 ng) dose of 5Z-7-oxozeaenol was given to Sprague-Dawley OIR rats at P14, followed by quantitative analysis of retinal neovascularization of flat mounted retinae at P18. (B) Retinal neovascularization is highlighted in white, and insets show selected areas at high magnification. Arrowheads indicate vessel tufts. Retinal neovascularization was quantified from 10 individual retinas per group. Scale bar: 500 µm. (C) Vaso-obliteration was simultaneously evaluated with the same samples analyzed in Fig. 3B. The white areas represent the avascular area in the retina. No significant difference was observed between groups (n = 10). Scale bar: 1 mm. (D) Vessel area of superficial and deep vascular layers was quantified in P18 control (n = 4) and OIR rats following a single intravitreal injection of vehicle, low or high dose of 5Z-7-oxozeaenol at P14 (n = 5 each group). Scale bar: 100 µm. Group data are shown as means ± SEM. Statistical analysis was undertaken with one-way ANOVA and Tukey’s multiple comparison test; \*\*\*\**P* < 0.0001.

### TAK1 deletion impairs the expression of multiple cytokines and cell adhesion molecules in endothelial cells responding to inflammatory insults

Given that TAK1 blockade prevented retinal neovascularization and previous studies have indicated that TAK1 is activated in response to various pro-inflammatory stimuli such as TNFα, IL-1β, TGF-β, or TLR ligand, we evaluated what signaling regulates TAK1 in the endothelial response to inflammation. We first generated TAK1-knockout (KO) human telomerase-immortalized microvascular endothelial (TIME) cells (TAK1-KO) using CRISPR/Cas9 (**Fig. S7**). To explore the effect of TAK1 deletion in human microvascular endothelial cells, we generated a transcriptome profile through bulk-RNA sequencing of wild-ype TIME cells (WT) and TAK1-KO TIME cells (TAK1-KO). We found that 2178 genes (743 upregulation and 1435 downregulation) were differentially expressed [false discovery rate (FDR), <0.05] following TAK1-KO (**Fig. S8A and S8B**). GSEA results suggested that TAK1 knockout negatively affected pathways related to tissue immune responses, cell adhesion, and cell-extracellular matrix interaction (**Fig. S8C**).

Since TNFα is strongly correlated with progression of retinal neovascularization and is a major activator of TAK1 signaling (10), we performed bulk-RNA sequencing to obtain transcriptomes from wild-type (WT) TIME cells and TAK1-KO TIME cells that were in the presence or absence of TNFα (**Fig. 4A**). We found that 462 and 3609 genes were differentially expressed (FDR<0.05 and Log2FoldChange >1 or <-1) in wild-type cells upon TNFα treatment (WT TNF *vs.* WT; hereafter referred to as the WT TNF set) and TAK1-KO cells upon TNFα treatment (TAK1-KO TNF *vs.* WT TNF; hereafter referred to as the TAK1-KO TNF set), respectively (**Fig. 4B**). To further explore the biological significance of our findings in differential gene expression, we undertook GSEA of WT TNF and TAK1-KO TNF sets. GSEA results showed the top 20 KEGG (Kyoto Encyclopedia of Genes and Genomes) gene sets were positively enriched due to TNFα stimulation in WT cells (**Fig. 4C**).

**Fig. 4.**
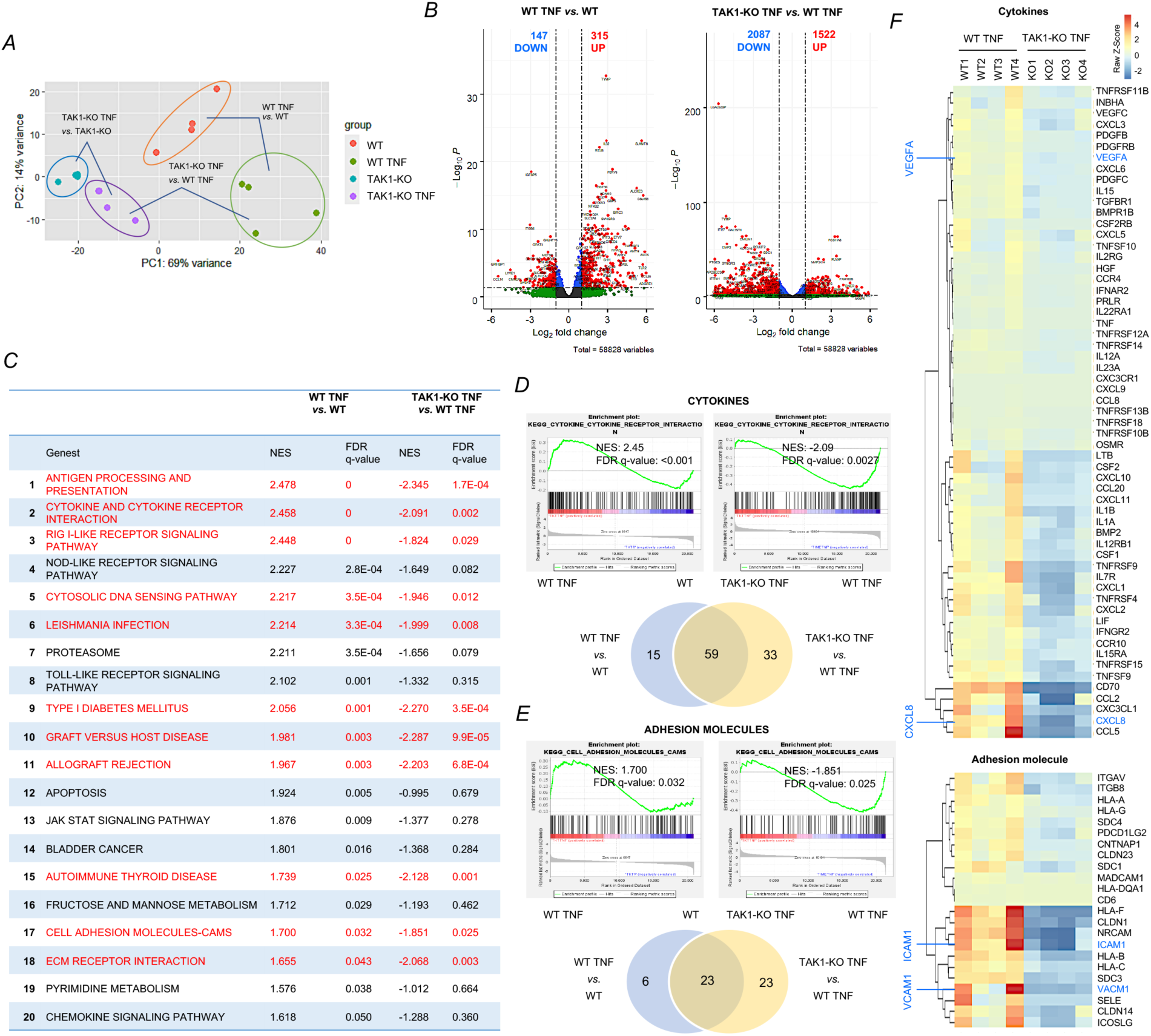
TAK1 deletion inhibits the expression of cytokines and cell adhesion molecules induced by TNFα in human microvascular endothelial cells. (A) Principal components analysis of RNA-seq data of wild-type (WT) endothelial cells (TIMEs), WT TIMEs treated with TNF*α*, TAK1 knockout (TAK1-KO) TIMEs, and TAK1-KO TIMEs treated with TNF*α*. Each dot represents an experimental replicate. (A) Volcano plots show genes that were significantly changed in both WT and TAK1-KO TIMEs when stimulated by TNF*α*. Red dots refer to genes showing a log2|fold change| > 1 and FDR < 0.05. Labels indicate significantly up-regulated genes in each data set. (C) GSEA of the top 20 KEGG gene sets in the two groups. Significantly enriched gene sets in WT TNF set (WT TNF vs. WT) (positive NES) and less enriched ones in TAK1-KO set (TAK1-KO TNF vs. WT TNF) (negative NES) are highlighted in red. False discovery rate (FDR) q-value < 0.05 was considered to be statistically significant. (D) GSEA plots for gene sets with Gene Ontology (GO) term “CYTOKINE AND CYTOKINE RECEPTOR INTERACTION” (related to cytokines) in the two groups. The Venn diagram shows the number of genes from the cytokine gene sets in the respective and overlapping areas of the two groups. (E) GSEA plots for gene sets with GO term “CELL ADHESION MOLECULES-CAMS” (related to adhesion molecules) in the two groups. The Venn diagram shows the number of genes from the chemokine gene sets in respective and overlapping areas of the two groups. (F) Heat map generated from the identified overlapping areas of “cytokines (n = 59)” “adhesion molecules (n = 23)” gene sets based on RNA-Seq read counts.

However, in TAK1-KO cells exposed to TNFα, 7 of the top 10 gene sets were negatively enriched, suggesting that these biological functions (highlighted in red in **Fig.4C**) induced by TNFα were adversely affected in cells due to TAK1 knockout. The gene sets related to cytokines (“Cytokine-cytokine receptor interaction”) and cell adhesion molecules (“Cell adhesion molecules-CAM”) were positively enriched in the wild-type cells exposed to TNFα yet not in TAK1-KO cells exposed to TNFα (**Fig. 4C**), indicating the expression of cytokines and cell adhesion molecules induced by TNFα were negatively affected in human microvascular endothelial cells due to TAK1 knockout. We further analyzed the target genes of TAK1 in these two gene sets by a Venn diagram. A total of 74 cytokine-related genes and 29 cell adhesion molecule-related genes were positively enriched in TNFα-treated WT cells (**Fig. 4D and 4E**). Among of these genes, 59 cytokine-related genes and 23 cell adhesion molecule-related genes were negatively enriched in TNFα-treated TAK1-KO cells (**Fig. 4D and 4E**). Many of these genes are related to inflammation and angiogenesis, suggesting that TAK1 may be involved in the regulation of the production of proinflammatory/proangiogenic cytokines (such as *CXCL8* and *VEGFA*), as well as leukocyte-endothelial interactions (leukostasis; such as *ICAM-1* and *VCAM-1*) in retinal neovascularization (**Fig. 4F**).

### TAK1 is required for activation of NFκB signaling in mediating the expression of multiple cytokines and cell adhesion molecules in endothelial cells responding to inflammatory insults

NFκB and MAPKs signaling are the primary known downstream pathways mediated by TAK1 activation, both of which have been shown as key mediators involved in several pathological conditions associated to angiogenesis, such as arthritis and cancer (25, 26) (**Fig. 5A**). We therefore investigated whether the identified 59 cytokine-related genes and 23 cell adhesion molecule-related genes from TNFα-treated TAK1-KO cells are regulated by NFκB and MAPKs signaling pathway. We found that about 75% (38/59) of selected genes from the “cytokine gene set” and 35% (8/23) of selected genes from the “adhesion molecule gene set” are known to be regulated by NF*κ*B and activator protein-1 (AP-1, a transcription factor regulates by MAPK signaling cascade), both of which were identified from a NF*κ*B target gene database (27) and ENCODE Transcription Factor Targets Database, respectively (**Fig 5B and Table 1**).

**Fig. 5.**
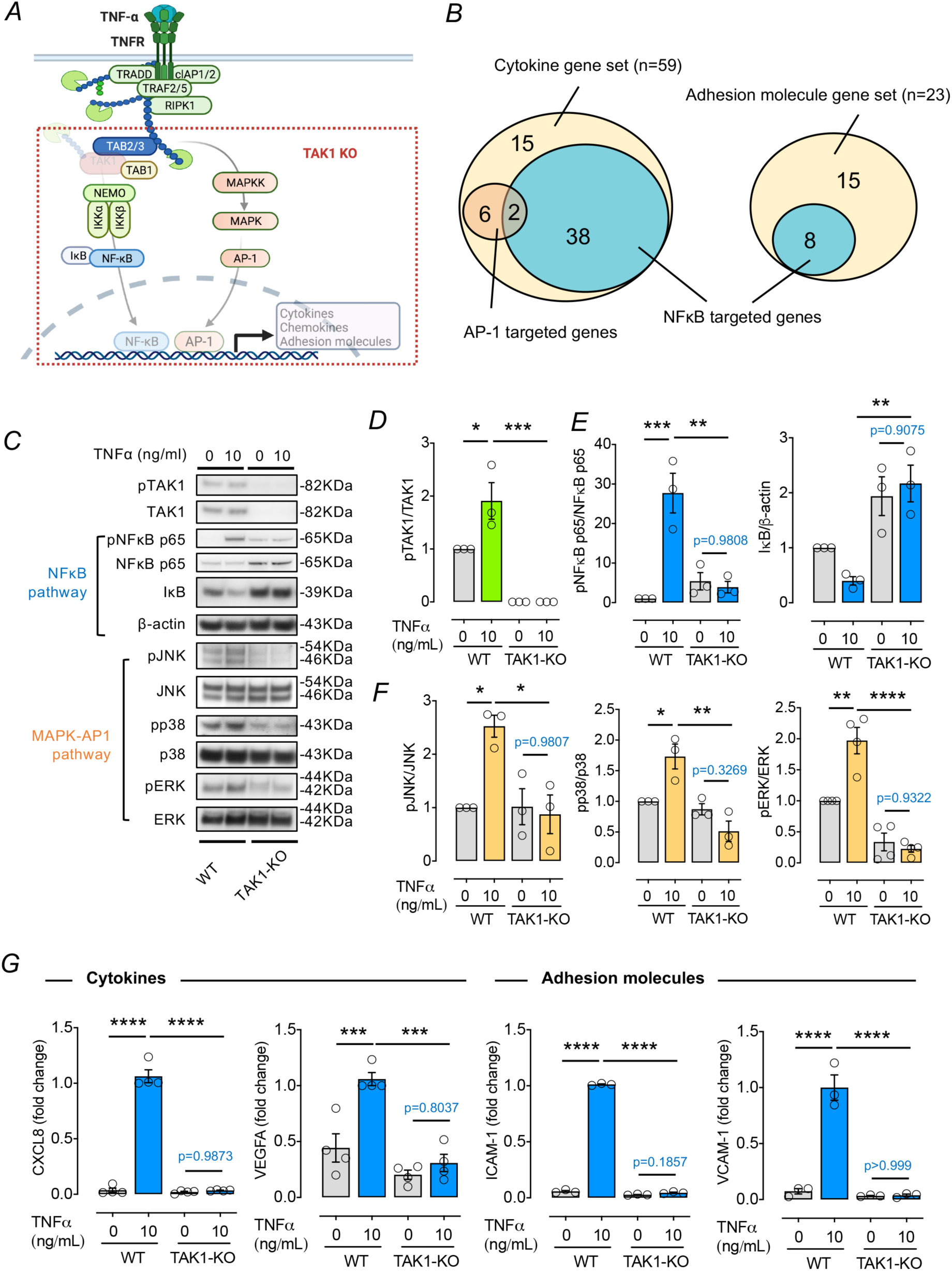
TAK1 deletion prevents TNFα-induced inflammatory cascade through the NFκB and MAPK pathway in human microvascular endothelial cells. (A) Schematic diagram of TNFα-TAK1 signaling. Upon TNFα stimulation, TAK1 binds to the polyubiquitin chain though TAB2, and activates the IKK complex, leading to the activation of NF-κB. TAK1 also activates MAPKs cascades. NFκB and MAPKs/AP-1 induce expression of inflammatory cytokines and cell adhesion proteins. (B) The Venn diagram respectively shows the number of genes from the “cytokine” and “adhesion molecule” gene sets, and the overlap indicates the number of genes directly targeted by NFκB or AP-1. (C) Western blot characterization of the TNFα-induced phosphorylation of TAK1, NFκB, and MAPKs proteins in TAK1 knockout (TAK1-KO) TIMEs. Wild-type (WT) and TAK1-KO TIMEs were stimulated by TNFα (10 ng/ml) for 10 minutes. Activation of TAK1, NFκB p65, IκB, JNK, p38, and ERK assessed using western blot (n = 3-4). Genetic TAK1 deletion reduced TAK1 expression (D) and impaired NFκB (NFκB p65 and IκB) (E) and MAPK (p38, JNK, and ERK) (F) signaling in TIMEs treated with TNFα. (G) qPCR results revealed that TAK1 deletion suppressed the expression of selected cytokines (*CXCL8* and *VEGFA)* and cell adhesion molecules (*ICAM-1* and *VCAM-1)* in TIMEs stimulated with TNFα for 24 hours (n = 3-4). Group data are shown as means ± SEM. Statistical analysis was undertaken with one-way ANOVA and Tukey’s multiple comparison test; \**P* < 0.05, \*\**P* < 0.01, \*\*\**P* < 0.001, \*\*\*\**P* < 0.0001.

To confirm the impairment of the activation of NFκB and MAPKs signaling caused by TAK1 knockout, we characterized the activation of NFκB and MAPKs in TNFα-treated TAK1-KO cells. TNFα treatment significantly increased TAK1 phosphorylation in WT cells, an effect not observed in TAK1-KO cells (**Fig. 5C and 5D**). IκB degradation and NFκB p65 phosphorylation are key events in NFκB signaling (28). TNFα treatment significantly induced NFκB p65 phosphorylation and IκB degradation in WT cells, while no such changes occurred in TAK1-KO cells (**Fig. 5C and 5E**). Likewise, phosphorylation of JNK, p38, and ERK in MAPK signaling was not increased in TNFα-treated TAK1-KO cells compared to TNFα-treated wild-type cells (**Fig. 5C and 5F**). Collectively, these data indicate that TAK1 is required for phosphorylation of the key mediators of NFκB and MAPK signaling in human microvascular endothelial cells.

Expression of the most significant genes (those also shown to be involved in retinal neovascularization) from the gene sets related to cytokines (“Cytokine-cytokine receptor interaction”) and cell adhesion molecules (“Cell adhesion molecules-CAM”) was also validated using qRT-PCR. Upon TNFα stimulation, the expression level of *CXCL8*, *VEGF-A, ICAM-1*, and *VCAM-1*, were substantially elevated in WT cells, while no TNFα induced change was observed in TAK1-KO cells (**Fig. 5G**). Similar results were found in IL1*β*-treated WT cells but not in IL1*β*-treated TAK1-KO cells (**Fig. S9**). Likewise, TAK1 knockdown using TAK1 siRNA (siTAK1) or TAK1 inhibitor (5Z-7-oxozeaenol) resulted in a similar effect in primary human retinal microvascular endothelial cells (HRMECs) (**Fig. S10 and S11**). These data further confirmed that TAK1 is required for activation of NFκB/AMPKs signaling to regulate expression of multiple cytokines and cell adhesion molecules in human microvascular endothelial cells responding to inflammation.

### Pharmacological inhibition of TAK1 suppresses the expression of inflammatory cytokines and reduces leukocyte-endothelial interactions (leukostasis) in the rat OIR model, while has no impact on retinal structure and function

To further confirm the effect of TAK1 inhibition in suppressing the expression of inflammatory cytokines and reduces leukostasis in the development of retinal neovascularization, OIR rats were treated with 5Z-7-oxozeaenol at P14 and the retinae were harvested for qPCR and histological analysis at the time points indicated in the figure (**Fig. 6A**). Two days after drug administration, TAK1 inhibition attenuated the expression of inflammatory and angiogenic genes, including *TNFα*, *VEGFA* and *ICAM-1*, in the retina of OIR rats that had received low (18 ng) and high doses (90 ng) of 5Z-7-oxozeaenol, respectively, compared to OIR rats that had vehicle only, suggesting that 5Z-7-oxozeaenol can mitigate the production of inflammatory and angiogenic cytokines in the progression of retinal neovascularization (**Fig. 6B**). Since TAK1 was found to modulate the expression of cell adhesion molecules in human endothelial cells responding to proinflammatory cytokine, we therefore further evaluated the event of microglial adhesion to the vascular surface (leukostasis). Co-immunostaining of isolectin GS-IB4 (blood vessel) and Iba1 (a marker of microglia) illustrated that 5Z-7-oxozeaenol treatment at both low and high doses significantly attenuated the microglial adhesion to the vascular surface in OIR rats (**Fig. 6C and 6D**). Together, our data suggest that active involvement of TAK1 in the regulation of proinflammatory and proangiogenic cytokine expression as well as modulation of leukostasis in the retinal vasculature. Thus, 5Z-7-oxozeaenol is therapeutically effective to suppress retinal neovascularization in the rat model of retinopathy (**Fig. 6E**).

**Fig. 6.**
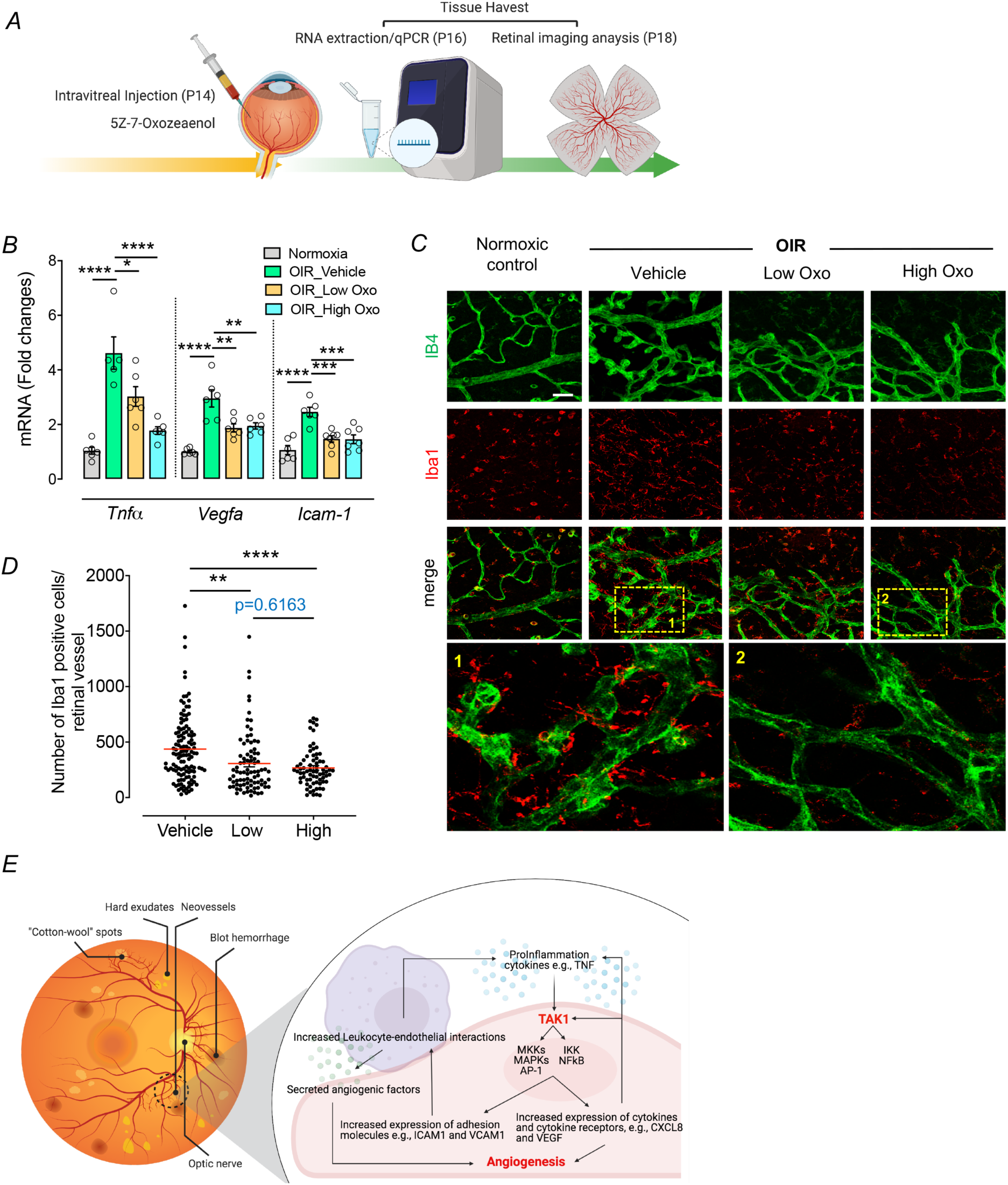
TAK1 inhibition by 5Z-7-oxozeaenol decreased inflammatory gene expression and adhesion of microglia to vessels in OIR. (A) Experimental protocol for 5Z-7-oxozeaenol treatment in the OIR rat model. A single intravitreal injection of vehicle, low (18 ng) or high (90 ng) dose of 5Z-7-oxozeaenol was given to Sprague-Dawley OIR rats at P14, and retinae were collected for qPCR and immunostaining at P16 and P18. (B) Intravitreal injection of 5Z-7-oxozeaenol in OIR rats at P14 inhibited *TNFα*, *VEGFA* and *ICAM-1* gene expression assessed using qPCR (n = 5-7). (C) Representative images of microglial adhesion to the vascular surface in a retina from a control rat and OIR rats that had a single intravitreal injection of vehicle, low or high dose of 5Z-7-oxozeaenol (Oxo). Vessels were visualized with isolectin GS-IB4 staining (green), and distribution of microglia was immunolabeled with Iba-1 (red). Scale bar: 50 µm. (D) Quantitative analysis of microglial number relative to retinal vessel area in vehicle or 5Z-7-oxozeaenol treated OIR rats (n = 7-10; 10 images from each retina were shown as separated dot plot). (E) Schematic diagram for the role of TAK1 signaling in microvascular endothelial cells. Proinflammatory cytokines such as TNFα can activate TAK1 in microvascular endothelial cells, and TAK1/TAB1/TAB2-3 complex can lead to activation of NFκB and MAPKs/AP-1 cascades. NFκB and MAPKs/AP-1 induce expression of “cytokines” and “adhesion proteins”, thereby triggering the “inflammatory cycle” and “leukostasis”, and leading to retinal neovascularization. Group data are shown as means ± SEM. Statistical analysis was undertaken with one-way ANOVA and Tukey’s multiple comparison test; \**P* < 0.05, \*\**P* < 0.01, \*\*\**P* < 0.001, \*\*\*\**P* < 0.0001.

In addition to examining the therapeutic efficacy of 5Z-7-oxozeaenol in retinal neovascularization, we further performed safety evaluations to test whether 5Z-7-oxozeaenol adversely affects rodent retinae. Electroretinography (ERG) and optical coherence tomography (OCT) were performed to look for any retinal functional or structural changes in rats 28 days post intravitreal injection of high dose of 5Z-7-oxozeaenol. No difference in function was observed for photoreceptors, bipolar cells or ganglion cells between vehicle- or 5Z-7-oxozeaenol-treated eyes (**Fig. S13A-S13F**). No significant difference was observed in total retinal thickness or thickness of various retinal layers including the nerve fiber layer (NFL) and outer retina layer (ORL) between 5Z-7-oxozeaenol- and vehicle-treated eyes (**Fig. S13G and S13H**). Together, these results suggested that intravitreal injection of 5Z-7-oxozeaenol does not result in adverse or toxic effects on retinal function and structure up to 28 days post-injection.

## DISCUSSION

TAK1 is a key modulator involved in a range of cellular functions including the immune response (14), cell survival and death (16), angiogenesis (20), fibrosis (29) and tumor metastasis (30). TAK1 is activated in response to various stimuli such as pro-inflammatory cytokines, hypoxia, oxidative stress, metabolism, DNA damage, Wnt and osmotic shock (14, 31–33). Whether TAK1 is involved in angiogenesis or inflammatory signaling in the retina, much less the molecular mechanisms by which this occurs, has not been studied. Here, we show that TAK1 plays a central role in regulating inflammatory and angiogenic signaling in the rat retina, and that the same signaling appears to be involved in human diabetic retinopathy. Pharmacological antagonism of TAK1 signaling by 5Z-7-oxozeaenol was shown to be a potential therapeutic option for retinal neovascularization.

NFκB and MAPK are critical regulators of stress responses, immunity, inflammation and angiogenesis. A variety of proinflammatory cytokines such as TNFα and IL1*β* activate these signaling pathways through a common upstream kinase TAK1 (34). For instance, TNFα binds to its receptor 1 (TNFR1) upon inflammatory stimuli, triggering TAK1 that in complex with TAB1-3 phosphorylate IKK at Ser^177^ and Ser^181^, which in turn leads to rapid proteasome-mediated degradation of the I*κ*B subunit. Subsequently, NFκB (p50/p65) is activated and translocated to the nucleus to promote gene transcription (17). In addition, the TAK1 complex phosphorylates MAPKs to activate the JNK pathway and the p38 MAPK pathways (both mediated by MKK4 and MKK7) as well as ERK by an IKK2/tumor progression locus 2 (TPL2)-dependent pathway (35). Phosphorylation of p38 affects F-actin polymerization in vascular endothelial cells and endothelial migration (36). Activation of JNK and ERK subsequent to p21-activated kinase 1 stimulation promotes endothelial migration through enhancement of cell motility (37). In particular, ERK is involved in VEGF-induced endothelial proliferation, migration (via actin polymerization, lamellipodia and filopodia formation and myosin contraction), morphogenesis (tube formation) and extracellular matrix degradation during angiogenesis (37). In fact, deletion of TAK1 leads to the inactivation of both JNK and NFκB signaling, as well as downregulation of expression of pro-survival genes in mice (38). Hence, deletion of TAK1 causes bone marrow and liver failure in mice due to the massive apoptotic death of hematopoietic cells and hepatocytes, ultimately resulting in death within a few days. Moreover, endothelial-specific TAK1 knockout mice demonstrated intestinal and liver hemorrhage secondary to apoptosis of endothelial cells and damaged vessels, resulting in rapid death (19). These data suggest that TAK1 protects endothelial cells and maintains vascular integrity and homeostasis under inflammatory conditions.

Here, we provide the first evidence that both genetic and pharmacological inhibition of TAK1 activity in human microvascular endothelial cells prevents inflammatory stimuli from inducing sequential phosphorylation of upstream kinases in both NFκB (p65) and MAPKs (JNK, p38 and ERK) pathways and suppressed expression of downstream genes that drive inflammation and angiogenesis. GSEA results confirmed that TAK1 knockout prevented TNFα induced cytokines and cell adhesion molecules expression predominantly through the NFκB pathway, resulting in attenuated inflammation and angiogenesis. In addition, TAK1 inhibition by 5Z-7-oxozeaenol in OIR rat significantly reduced cytokines expression and adhesion of microglia to retinal endothelial cells. These data together suggest a significant role for TAK1 in mediating crosstalk between inflammatory and angiogenic pathways, both of which are crucial in pathological angiogenesis.

TAK1 is thus a central regulator of cell survival and death (16). 5Z-7-oxozeaenol, a resorcylic acid lactone derived from a fungus, acts as a potent and selective inhibitor of TAK1, which displays more than 33- and 62-fold selectivity for TAK1 over binding of MEKK1 and MEKK4, respectively (23). It forms a covalent complex with TAK1 and impedes both the kinase and ATPase activity of TAK1 following bi-phase kinetics (39). We demonstrated that intravitreal injection of 5Z-7-oxozeaenol suppressed retinal neovascularization, and modulated elevated TNFα expression in the retina of OIR rats, without causing degeneration of the normal vasculature. These data suggest that TAK1 inhibition by 5Z-7-oxozeaenol may more specifically attenuate active angiogenesis with less risk of damage to normal endothelial cells. However, further studies are required to understand the effects of repeated 5Z-7-oxozeaenol treatment. Moreover, TAK1 Inhibition by 5Z-7-oxozeaenol is known to block pro-inflammatory signaling by selectively inhibiting TAK1 MAPKKK (23). In an animal model of early brain injury, 5Z-7-oxozeaenol was shown to inhibit subarachnoid hemorrhage-induced phosphorylation of p38 and JNK, the nuclear translocation of NFκB p65 and degradation of IκB, ultimately reducing neuronal apoptosis and early brain injury (40). Similarly, another study demonstrated that 5Z-7-oxozeaenol alleviated experimental autoimmune encephalomyelitis by reducing the levels of pro-inflammatory cytokines in splenocytes and the central nervous system, thus reducing the number of activated microglia and blocking p38 MAPK, JNK, and ERK signaling pathways (41). Overactive microglia can damage surrounding cells, via a range of detrimental effects including release of pro-inflammatory cytokines, reactive nitrogen species, reactive oxygen, and proteolytic enzymes (42). We also observed diminished adhesion of microglia to endothelial cells in OIR rats receiving 5Z-7-oxozeaenol treatment, suggesting that 5Z-7-oxozeaenol alleviates inflammation in the retina, in part by reducing activation of microglia. Altogether, our results indicate that TAK1 inhibition by 5Z-7-oxozeaenol suppresses ocular inflammation and neovascularization by inhibiting cellular proliferation and microglia activation through NFκB pathway.

5Z-7-oxozeaenol has been previously examined in a number of animal studies focusing on brain injury (43–45). One study demonstrated that 5Z-7-oxozeaenol has neuroprotective function in a mouse model of stroke (43). However, others suggest long-term TAK1 inhibition may not be protective in neurodegenerative disorders (46). Chronic TAK1 inhibition resulted in apoptosis of a number of normal cells, including keratinocytes, hepatocytes, and hematopoietic cells, raising safety concerns (38, 47, 48). We demonstrated that TAK1 inhibition was capable of simultaneously reducing proinflammatory cytokines and proangiogenic cytokines in the retina of OIR rats. In our studies, no adverse effects of intravitreal injection of the TAK1 inhibitor on retinal function or structure were apparent up to 28 days post-injection. Specifically, TAK1 inhibition by intravitreal injection of 5Z-7-oxozeaenol did not affect the scotopic threshold response, a highly sensitive electroretinography measure of retinal ganglion cell activity (49). Another study has shown that systemic TAK1 inhibition by 5Z-7-oxozeaenol preserved β cell function, thus improving overall health condition in the animal with autoimmune diabetes (50). It is worth noting that 5Z-7-oxozeaenol may have off-target effects, potentially stemming from cross-reactivity with other human kinases that possess an analogous cysteine (51). Whilst encouraging, further studies with chronic dosing of 5Z-7-oxozeaenol are clearly needed to confirm safety.

In summary, the present study identifies TAK1 as a mediator of inflammatory and angiogenic signaling pathways, with a central role in retinal neovascularization. We demonstrated elevated TAK1 signaling in both human and rat retinae associated with neovascularization. We showed that reducing TAK1 suppressed the expression of multiple cytokines and leukostasis in endothelial cells responding to inflammatory insults *in vitro* through NFκB signaling pathway. Using 5Z-7-oxozeaenol, a well-studied selective TAK1 inhibitor, we demonstrated that pharmacological inhibition of TAK1 can effectively suppress retinal neovascularization in *in vivo* models in rats. These findings provide new insights into the molecular and cellular mechanisms by which TAK1 modulates pathological angiogenesis, thus highlighting a potential alternative approach to anti-VEGF treatment to prevent retinal neovascularization.

## MATERIALS AND METHODS

### Data accession

To assess TAK1 expression in human retinal cells, we utilized data single cell transcriptomes from human retina, from the Gene Expression Omnibus (https://www.ncbi.nlm.nih.gov/geo/query/acc.cgi?acc=GSE135133) (accessed on September 20, 2020).

To access gene expression in retinas from patients with retinal neovascularization due to proliferative diabetic retinopathy, we obtained the RNA-seq raw counts and TPM matrices from the Gene Expression Omnibus (https://www.ncbi.nlm.nih.gov/geo/query/acc.cgi?acc=GSE102485) (accessed on September 27, 2020). We specifically compared samples from patients with retinal neovascular proliferative membranes due to type II diabetes (GSM2739349, GSM2739350 and GSM2739351) and those from normal controls (GSM2739364, GSM2739365 and GSM2739366).

### scRNA-seq data analysis

scRNA-seq data were retrieved from a public dataset (GSE135133) and analyzed in R (v3.6.3) using Seurat (v3.2) (52, 53) with customized scripts. In brief, cell identity was assigned according to the assignment labels kindly provided by the authors after we normalized and scaled all human samples. We assigned ganglion cells, horizontal cells, bipolar cells, amacrine cells, cones and rods as the neural cells while immune cells, basal cells, red cells, pericytes, endothelial cells, astrocytes and Müller cells were regarded as non-neural cells. Dot plots were utilized for visualization of gene expression levels and percentage difference for various cell types.

### RNA isolation

Total cells or tissue RNA was isolated using Zymo Quick-RNA MiniPrep kit (R1055; Zymo Research, Irvine, CA) according to the manufacturer’s instructions. The quantity of RNA was evaluated with a NanoDrop ND-1000 spectrophotometer (NanoDrop Technologies, Wilmington, DE).

### RNA-seq analysis

Analysis was undertaken in triplicate on total RNAs (1 μg) from normoxic and OIR rats at P14, prepared as per the manufacturers’ instructions. cDNA libraries were sequenced by using an Illumina Hiseq-2500 RNA-seq platform as 50 bp single end chemistry at the Australian Genome Research Facility (AGRF, Melbourne, VIC, Australia). The raw fastq files of single-end reads were removed the adapter sequences and the low-quality reads were dropped by Trimgalore 0.4.4 (Babraham Bioinformatics). Filtering parameter were set as follow: -q 25 --length 50 -e 0.1 --stringency 5. The trimmed reads were subjected to alignment with default setting using STAR 2.5.3. The reference genomes for human and rat were hg38 and rn6, respectively. Aligned RNA-seq data was counted over gene exon using feature Counts (subread 1.6.4). Genes were annotated as per the Gencode Version 33 annotation file.

### Gene set enrichment analysis (GSEA)

GSEA was run on the human dataset of protein-coding genes in PDR versus controls and on the rodent dataset of protein-coding genes in OIR versus controls, against canonical pathway genesets collection (c2.cp.v7.2, Broad Institute) by using 1000 genesets permutations. Only genesets with FDR < 0.05 were considered. Readcount values were loaded into R (v3.6.3) for statistical analysis and the heatmap function was used to perform hierarchical clustering analysis and for data representation.

### Quantitative polymerase chain reaction (qPCR)

Total RNAs (100 ng) was reverse transcribed to cDNA using a High-Capacity cDNA Reverse Transcription kit (4368814; Life Technologies, Carlsbad, CA). Quantitative PCR was performed using TaqMan fast advanced master mix (4444553; Applied Biosystems, Foster City, CA) and gene TaqMan probe sets to detect gene expression (**Table S2**). The expression levels of target genes were normalized to the levels of rat beta-actin (Rn00821065_g1) or human GAPDH (Hs99999905_m1). Subsequently, the ΔΔCt method was used to evaluate relative expression level (fold change) in each condition versus the corresponding control condition.

### Cell culture

Telomerase-immortalized human microvascular endothelial cell (TIME; CRL-4025™) and primary human retinal microvascular endothelial cell (HRMEC; ACBRI 181V) were purchased from American Type Culture Collection (ATCC; Manassas, VA) and Cell Systems (Kirkland, WA). Endothelial cells were cultured in EBM^TM^ Plus Basal Medium supplied with EGM^TM^ Plus SingleQuots^TM^ supplements (CC-5035; Lonza, Basel, Switzerland). All cell lines were maintained in a humidified incubator at 37°C and 5% CO_2_. Cell lines were mycoplasma free tested by MycoAlert™ Mycoplasma Detection Kit (LT07; Lonza).

### Western blotting

A standard immunoblotting protocol was used. Briefly, protein was extracted from cells or retinas with Pierce RIPA buffer (89900; Life Technologies) supplied with protease inhibitor cocktail (11697498001; Roche Diagnostics, Basel, Switzerland). Equal amounts of cell lysates from different groups were subjected to Western blotting analysis with specific antibodies against IκB, phospho-NFκB p65, NFκB p65, phospho-p44/42 MAPK (Erk1/2), p44/42 MAPK (Erk1/2), phospho-TAK1, TAK1, phospho-SAPK/JNK, SAPK/JNK, phospho-p38 MAPK, p38 MAPK (4814, 3033, 4764, 9101, 4695, 9339, 4505, 9255, 9252, 9211, 8690, respectively; Cell Signaling), and β-actin (MAB1501; Merck Millipore, Burlington, MA). The quantitative densitometry was analyzed using Image Pro Plus software (Media Cybernetics, Rockville, MA).

### Animals

Pregnant Sprague-Dawley rats (female, 12-14 weeks old) and Brown Norway rats (male, 12 weeks old) were supplied by the Animal Resources Centre (Murdoch, WA, Australia) and Cambridge Farm Facility of the University of Tasmania (Cambridge, TAS, Australia). Animals were housed in standard cages with a temperature/humidity-controlled and a 12-hour light (50 lux illumination)/12-hour dark (<10 lux illumination) cycle environment. Food and water were available *ad libitum*. All animal experiments were conducted in accordance with guidelines of the ARVO Statement for the Use of Animals in Ophthalmic and Vision Research and were approved by the Animal Ethics Committees of St Vincent’s Hospital (004/16), the University of Melbourne (13-044UM), and the University of Tasmania (A0017311). Rats were randomly allocated to treatment groups such that littermates were distributed equally between groups.

### Rat model of oxygen-induced retinopathy and intravitreal injection of 5Z-7-oxozeaenol

The oxygen-induced retinopathy model was induced in rats as previously described (54). Briefly, Sprague-Dawley litters (within 12 hours of birth; P0) and their nursing mothers were exposed to daily cycles of 80% O_2_ for 21 hours and room air for 3 hours in a custom-built and humidity-controlled (< 80%) chamber until P14. At P14 animals were returned to room air until P16 or P18, when they were sacrificed to harvest retinae. An ProOx 110 oxygen controller (BioSpherix; Parish, NY) was used to regulate and monitor the oxygen level. At P14, intravitreal injections were performed to administer 5Z-7-oxozeaenol visualizing with a microsurgical microscope. Under surface anesthesia, a puncture in the superior temporal quadrant of the limbus was made by a 30-gauge needle. A hand-pulled glass micropipette (Bio-Strategy, Australia) connected to a 10 μl Hamilton syringe (Hamilton Company, Reno, NV) was inserted into the vitreous cavity, and one microliter of balanced salt solution containing 1% DMSO (as vehicle), low (18 ng) or high dose (90 ng) of 5Z-7-oxozeaenol was intravitreally injected into the eye. For safety evaluation, Brown Norway rats were intravitreally injected with 1 μl high dose (90 ng) of 5Z-7-oxozeaenol or vehicle. Eyes were excluded from the study if large backflow after injection or severe hemorrhage were occurred. Electroretinography and optical coherence tomography imaging evaluation was undertaken 28 days post-intravitreal injection in anesthetized animals as previously described (55).

### Immunohistochemistry and vessel quantification

Collection of donor samples was approved by the Human Research Ethics Committee of the Royal Victorian Eye and Ear Hospital (HREC13/1151H) and carried out in accordance with the approved guidelines. Post-mortem eye globes were obtained via the Lions Eye Donation Service (Royal Victorian Eye and Ear Hospital, East Melbourne, VIC, Australia) and processed for paraffin embedding and sectioning following standard histological procedures. Briefly, donated eyes were isolated, fixed, and permeabilized. The cross sections were stained with anti-TAK1 (1:100; NBP1-87819, Novus Biologicals, Littleton, CO) and anti-CD31 (1:25; M0823, DAKO, Jena, Germany) primary antibody and then with the appropriate secondary antibodies. Images were digitized using a confocal laser scanning microscope (FV3000; Olympus, Tokyo, Japan).

The rat retinal flat-mount were collected at P18 followed by retina dissection, staining overnight with isolectin GS-IB_4_ Alexa Fluor™ 488 conjugate (5 μg/mL; I21411, Life Technologies), anti-TAK1 (1:100; NBP1-87819, Novus Biologicals), or anti-Iba1 (1:300; 019-19741, FUJIFILM Wako Chemicals, Richmond, VA), and then incubated with the appropriate secondary antibodies. The images were captured by FV3000 confocal laser scanning microscope. The avascular (vaso-obliteration) and neovascularized areas were quantified by two blinded assessors using Adobe Photoshop (San Jose, CA) (56). Deep vessel area was analyzed by AngioTool software (National Institutes of Health, Bethesda, MA) (57).

### Statistical analysis

All data are expressed as mean ± SEM. Statistical analyses were performed with Graphpad Prism 7 (GraphPad, San Diego, CA) using student t-test for unpaired data, one-way or two-way ANOVA wherever appropriate followed by Tukey’s or Bonferroni’s multiple comparisons test. *P* values < 0.05 were considered statistically significant.

### Data and resource availability statement

The main datasets generated or analyzed during this study are included here in the article. Retinal RNA profiles of OIR and control rats have been deposited in Gene Expression Omnibus and are accessible through GEO Series accession number GSE104588 (https://www.ncbi.nlm.nih.gov/geo/query/acc.cgi?acc=GSE104588). The original Western Blot images and the details of statistical analysis for this study are available in the supplementary information. Other original datasets will also be available to the research community upon reasonable request.

## ACKNOWLEDGEMENTS

The authors thank UTAS CFF animal technicians, Karen Shiels, Keri Smith, Heather Howard, Lisa Harding and Danielle Eastley for their assistance with rat chamber operation. We also thank Dr Jacqueline Y.K. Leung for assistance with intravitreal injection, and Zheng He for assistance with retinal imaging. This work was supported by grants from the National Health and Medical Research Council of Australia (1061912, 1185600 and 1123329), the Ophthalmic Research Institute of Australia, the National Natural Science Foundation of China (82000902) and the Shenzhen Key Laboratory of Biomimetic Materials and Cellular Immunomodulation (ZDSYS20190902093409851). GJD was a Principal Research Fellow of NHMRC. The Centre for Eye Research Australia receives Operational Infrastructure Support from the Victorian Government.

## AUTHOR CONTRIBUTIONS

Conceptualization- F-L.L., J-H.W., P.vW., G.J.D., G-S.L. Methodology- F-L.L., C.M., J-H.W., B.V.B., P.vW., G-S.L. Formal Analysis- F-L.L., J-H.W., J.C., R.C.B.W., B.V.B., G-S.L. Investigation- F-L.L., J-H.W., J.C., L.Z., Y-F.C., L.T., C.M., S.L., D.L., C-L.T., B.V.B. Resources- R.C.B.W., A.W.H., C-L.T., G.J.D., P-Y.W., G-S.L. Data Curation- F-L.L., J-H.W., J.C., G-S.L. Writing (Original Draft)- F-L.L., J-H.W., G-S.L. Writing (Review & Editing)- J.C., C.M., R.C.B.W., B.V.B., P.vW., G.J.D., P-Y.W. Project Administration- F-L.L., J-H.W., G-S.L. Funding Acquisition- L.T., G.J.D., P-Y.W., G-S.L. Project Supervision- G.S.L.

## SUPPLEMENTARY MATERIALS

**Table S1.** Effects on the expression of enriched genes mediated by NFκB or AP-1 signaling in “TAK1-KO TNF *vs.* WT TNF” data set compared to “WT TNF *vs.* WT” data set.

| Gene name | WT TNF vs WT |  | TAK1-KO TNF vs TAK1-KO TNF |  | NFκB or AP-1 targeted gene |
| --- | --- | --- | --- | --- | --- |
|  | log2FoldChange | FDR q-value | log2FoldChange | FDR q-value |  |
| Cytokines-Cytokine Receptors (n=58) |  |  |  |  |  |
| IL1B | 4.35670 | 3.94E-11 | -4.41950 | 2.40E-05 | NFκB |
| CXCL1 | 1.83710 | 7.24E-11 | -2.34930 | 3.41E-04 | NFκB |
| IL1A | 3.29810 | 5.17E-10 | -1.87860 | 6.40E-03 | NFκB |
| VEGFC | 1.72730 | 1.06E-09 | -0.81200 | 1.29E-02 | NFκB |
| TNFRSF9 | 6.36650 | 1.62E-09 | -5.03510 | 2.97E-05 | None |
| TNFRSF4 | 4.86660 | 1.03E-08 | -2.56130 | 9.43E-03 | None |
| CCL2 | 4.79430 | 2.35E-08 | -3.10600 | 2.47E-02 | NFκB |
| CCL5 | 6.50100 | 1.47E-07 | -6.05740 | 5.44E-07 | NFκB |
| CXCL2 | 2.68970 | 8.93E-07 | -2.39200 | 1.56E-02 | NFκB |
| CD70 | 1.74470 | 1.94E-06 | -8.39220 | 4.56E-13 | NFκB |
| PDGFB | 1.05430 | 2.71E-06 | -0.63160 | 4.20E-04 | NFκB |
| LIF | 2.06620 | 8.24E-06 | -2.46700 | 2.83E-10 | NFκB |
| CXCL5 | 4.02040 | 9.25E-06 | -1.43094 | 3.26E-01 | None |
| CXCL6 | 3.20050 | 1.13E-05 | -3.24340 | 1.99E-05 | NFκB |
| CXCL8 | 3.40100 | 2.62E-05 | -4.62360 | 1.53E-07 | NFκB |
| CXCL3 | 2.83370 | 3.02E-05 | -1.95808 | 2.94E-01 | NA |
| CCR10 | 1.38350 | 5.53E-05 | -1.94630 | 6.67E-09 | AP-1 |
| CXCL10 | 9.88480 | 6.15E-05 | -5.92910 | 5.15E-03 | NFκB |
| CSF1 | 1.54050 | 9.12E-05 | -1.47410 | 2.91E-09 | NFκB |
| CXCL11 | 8.86230 | 1.07E-04 | -5.92440 | 9.63E-03 | NFκB |
| IL12RB1 | 2.38450 | 1.60E-04 | -2.56180 | 6.75E-08 | NFκB |
| IFNGR2 | 1.06720 | 2.21E-04 | -1.20550 | 3.02E-09 | NFκB |
| VEGFA | 0.97810 | 2.91E-04 | -0.96540 | 5.89E-07 | NFκB |
| CX3CL1 | 4.93410 | 3.00E-04 | -5.75310 | 1.09E-05 | NFκB |
| CCL20 | 5.98070 | 3.08E-04 | -6.66700 | 2.81E-04 | NFκB |
| TNFRSF11B | 2.27490 | 5.60E-04 | -1.22159 | 2.09E-01 | None |
| LTB | 5.02310 | 8.03E-04 | -5.28530 | 4.51E-07 | NFκB |
| IL15RA | 0.95000 | 1.28E-03 | -1.26430 | 8.62E-09 | NFκB |
| TNFSF9 | 1.00000 | 1.36E-03 | -2.03650 | 8.17E-15 | NFκB |
| TNFSF15 | 1.19150 | 2.94E-03 | -2.26180 | 4.67E-18 | None |
| **BMP2** | 1.54300 | 3.27E-03 | -1.69090 | 8.02E-03 | NFκB |
| **TGFBR1** | 0.70040 | 2.20E-02 | -0.82610 | 2.84E-05 | NFκB |
| **IL12A** | 1.68170 | 3.07E-02 | -1.53970 | 1.53E-02 | NFκB |
| **PRLR** | 2.91580 | 3.15E-02 | -3.27440 | 4.52E-03 | AP-1 |
| **IL15** | 0.85040 | 3.16E-02 | -1.26610 | 1.07E-06 | NFκB |
| **IL23A** | 1.82120 | 5.59E-02 | -1.58160 | 1.52E-02 | NFκB/AP-1 |
| **IL2RG** | 0.95541 | 1.01E-01 | -1.82390 | 5.10E-01 | None |
| **TNFRSF12A** | 0.71783 | 1.18E-01 | -0.43630 | 6.51E-03 | AP-1 |
| **TNFRSF14** | 0.46530 | 1.53E-01 | -0.48970 | 2.73E-03 | None |
| **TNFRSF10B** | 0.52018 | 1.97E-01 | -0.35370 | 4.89E-02 | NFκB |
| **BMPR1B** | 1.02127 | 1.98E-01 | -2.11260 | 2.99E-04 | None |
| **HGF** | 2.08440 | 2.92E-01 | -2.90270 | 1.53E-02 | NFκB |
| **PDGFC** | 0.42220 | 2.94E-01 | -0.84640 | 1.00E-05 | NFκB |
| **IL22RA1** | 3.47750 | 3.63E-01 | -5.93710 | 5.43E-03 | NFκB |
| **CCR4** | 2.87390 | 4.03E-01 | -3.34307 | 3.91E-01 | NA |
| **CSF2** | 1.64173 | 4.61E-01 | -0.17474 | 5.04E-01 | NFκB |
| **OSMR** | 0.40440 | 5.10E-01 | -0.24603 | 3.82E-01 | None |
| **TNFSF10** | 1.37807 | 5.88E-01 | -0.74565 | 3.64E-01 | None |
| **PDGFRB** | 0.42136 | 7.46E-01 | -1.75040 | 7.60E-05 | AP-1 |
| **TNFSF13B** | 3.28969 | NA | -3.31250 | 3.42E-02 | NFκB |
| **TNFRSF18** | 2.87284 | NA | -2.91772 | 1.48E-01 | AP-1 |
| **IL7R** | 3.67340 | NA | -3.50272 | NA | NFκB/AP-1 |
| **CX3CR1** | 2.38491 | NA | -2.44076 | NA | NFκB |
| **TNF** | 5.49810 | NA | -5.56714 | NA | NFκB |
| **CXCL9** | 3.32350 | NA | -3.39533 | NA | NFκB |
| **IFNAR2** | 0.34410 | NA | -0.39382 | 9.84E-02 | AP-1 |
| **INHBA** | 0.83100 | NA | -3.20137 | 2.55E-01 | None |
| **CCL8** | 2.91000 | NA | -2.98153 | NA | None |
| **CSF2RB** | 1.54048 | NA | -2.04434 | 6.12E-02 | None |
| **Cell Adhesion Molecules (n=23)** |  |  |  |  |  |
| **HLA-B** | 1.54750 | 1.24E-10 | -2.12490 | 4.67E-29 | NFκB |
| **NRCAM** | 2.08430 | 1.99E-10 | -3.42680 | 3.44E-08 | None |
| **CLDN14** | 2.72790 | 8.91E-10 | -1.73390 | 1.61E-02 | None |
| **HLA-F** | 2.57600 | 2.25E-07 | -4.58300 | 1.43E-14 | NFκB |
| **ICAM1** | 3.58140 | 4.18E-07 | -3.17350 | 3.58E-04 | NFκB |
| **VCAM1** | 10.39070 | 6.70E-07 | -7.44920 | 3.86E-04 | NFκB |
| **CNTNAP1** | 1.13470 | 8.81E-06 | -0.87440 | 3.03E-07 | None |
| ***ICOSLG*** | 3.32720 | 1.44E-05 | -3.15500 | 4.04E-03 | None |
| ***CLDN1*** | 2.09970 | 2.08E-05 | -4.77710 | 1.22E-12 | None |
| ***ITGAV*** | 1.70150 | 6.11E-05 | -1.20700 | 8.66E-03 | None |
| ***HLA-C*** | 0.85300 | 2.49E-04 | -1.31420 | 9.92E-15 | None |
| ***HLA-A*** | 0.83990 | 3.40E-04 | -0.81560 | 2.38E-06 | NFκB |
| ***SDC4*** | 0.91350 | 5.62E-04 | -0.79390 | 6.91E-05 | None |
| ***PDCD1LG2*** | 0.98570 | 2.49E-03 | -0.96010 | 1.02E-04 | NFκB |
| ***SDC1*** | 0.90460 | 1.09E-02 | -1.18610 | 1.06E-06 | None |
| ***SDC3*** | 0.41120 | 4.08E-02 | -1.86010 | 7.91E-51 | None |
| ***CLDN23*** | 1.91240 | 1.40E-01 | -4.50220 | 1.21E-05 | None |
| ***SELE*** | 4.44980 | 1.87E-01 | -9.34310 | 4.74E-05 | NFκB |
| ***ITGB8*** | 0.61910 | 2.80E-01 | -2.01250 | 1.03E-10 | NFκB |
| ***HLA-G*** | 0.77070 | 3.23E-01 | -1.45580 | 2.09E-03 | None |
| ***MADCAM1*** | 0.83220 | 7.39E-01 | -0.84027 | NA | None |
| ***HLA-DQA1*** | 2.94940 | NA | -3.01068 | NA | None |
| ***CD6*** | 1.94430 | NA | -2.00757 | NA | None |

FDR: False Discovery Rate, NA: Not Available, TAK1-KO: TAK1 knockout, TNF: TNFα , WT: Wild-type.

**Table S2.**
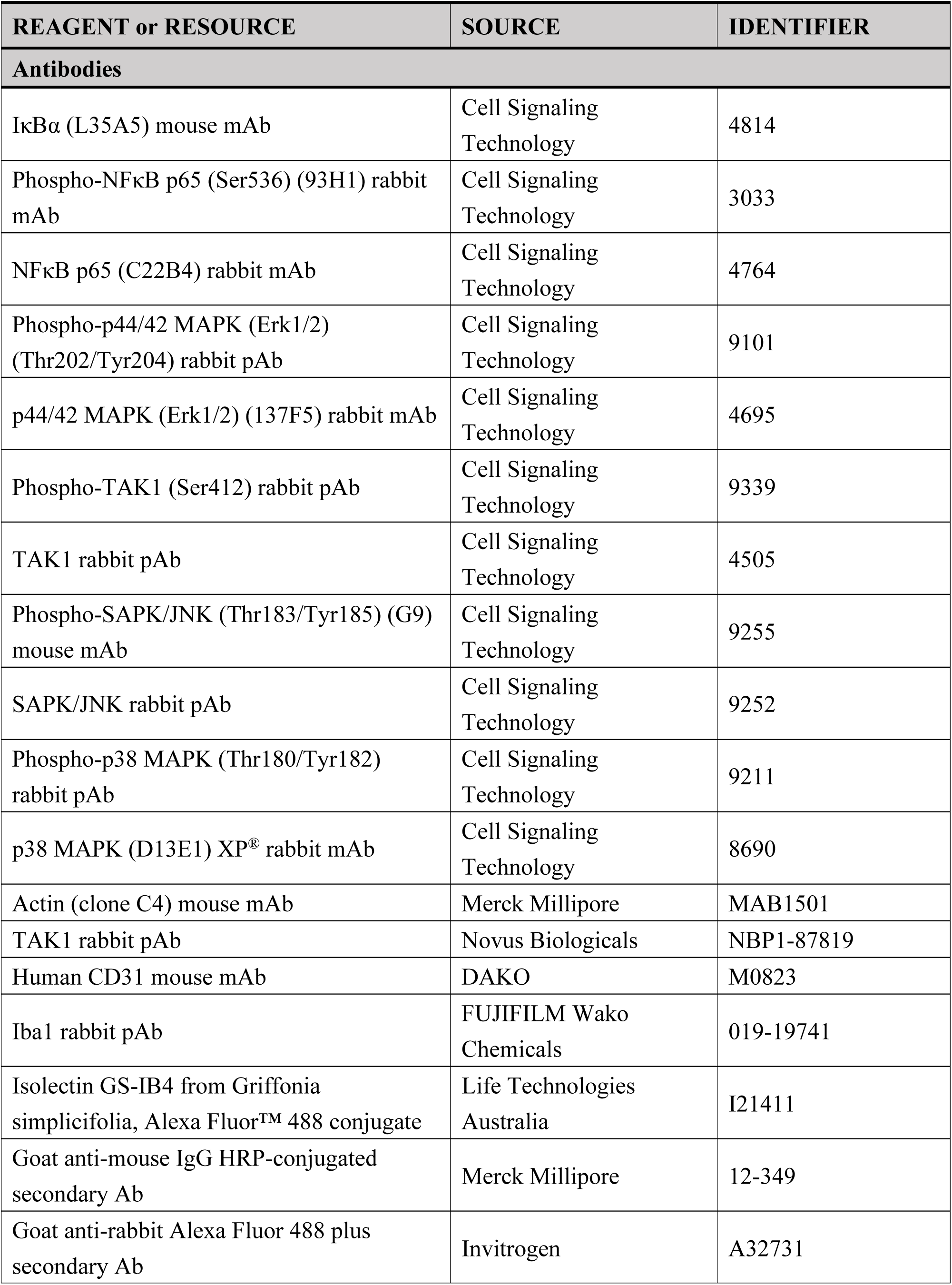
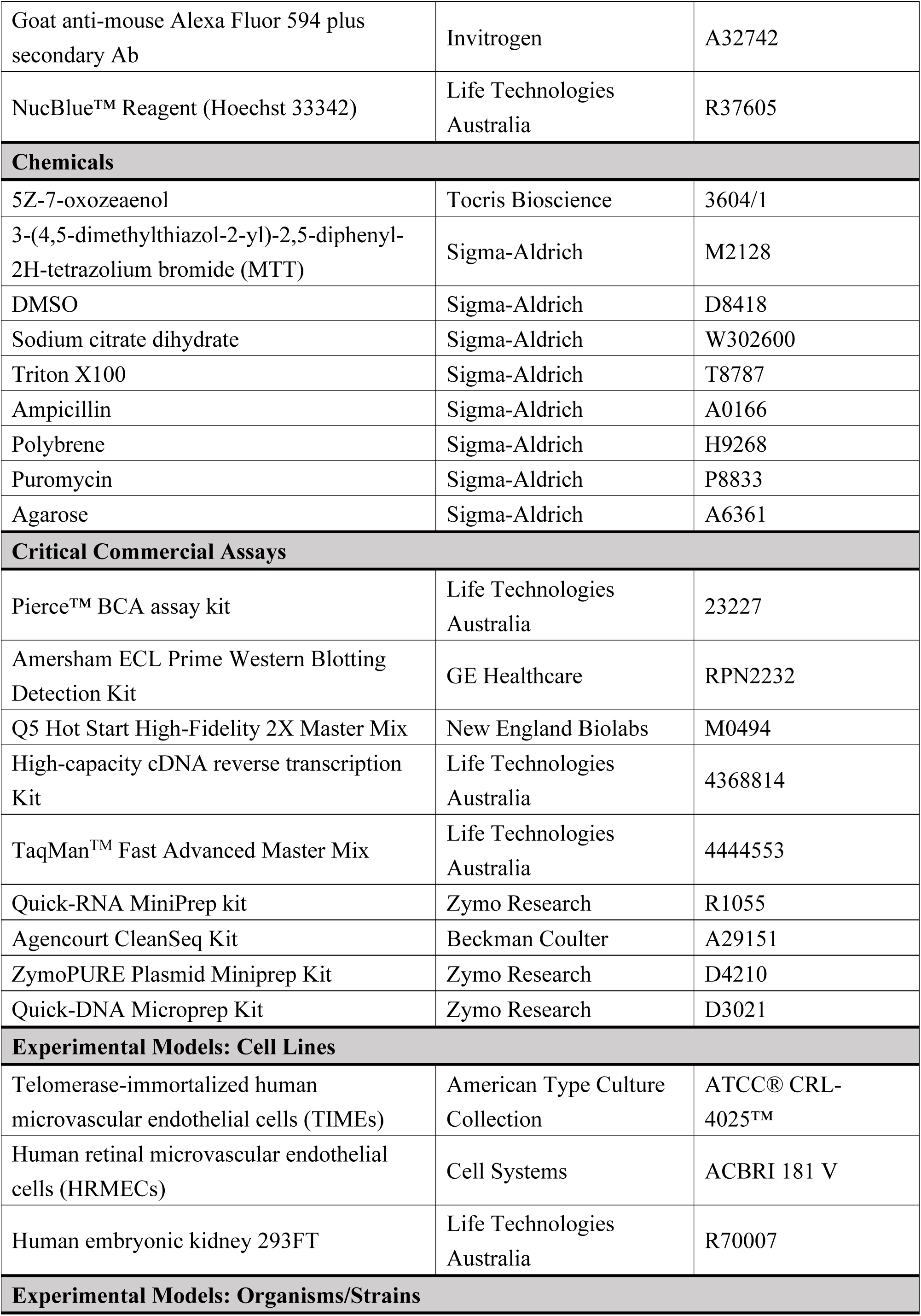
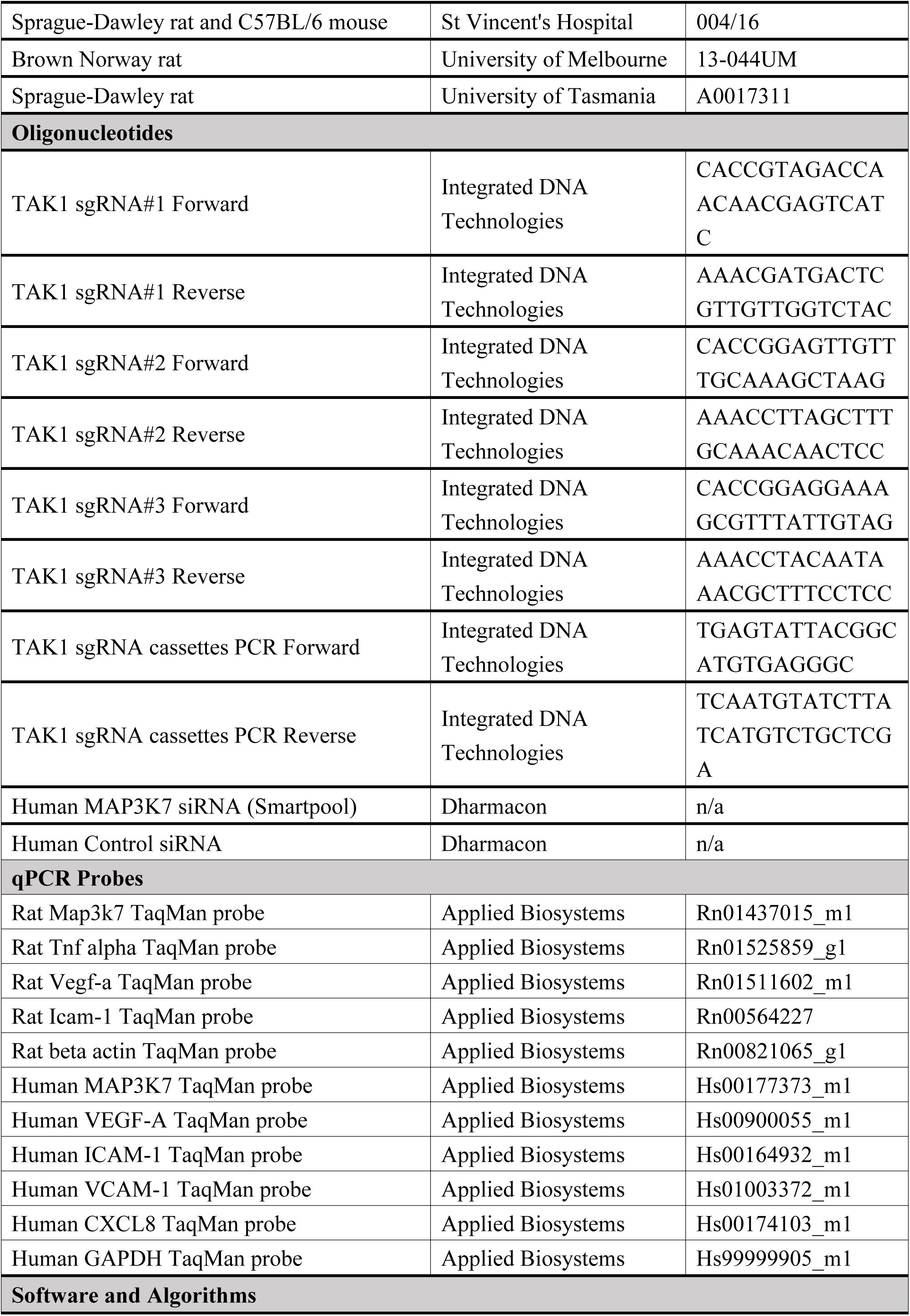
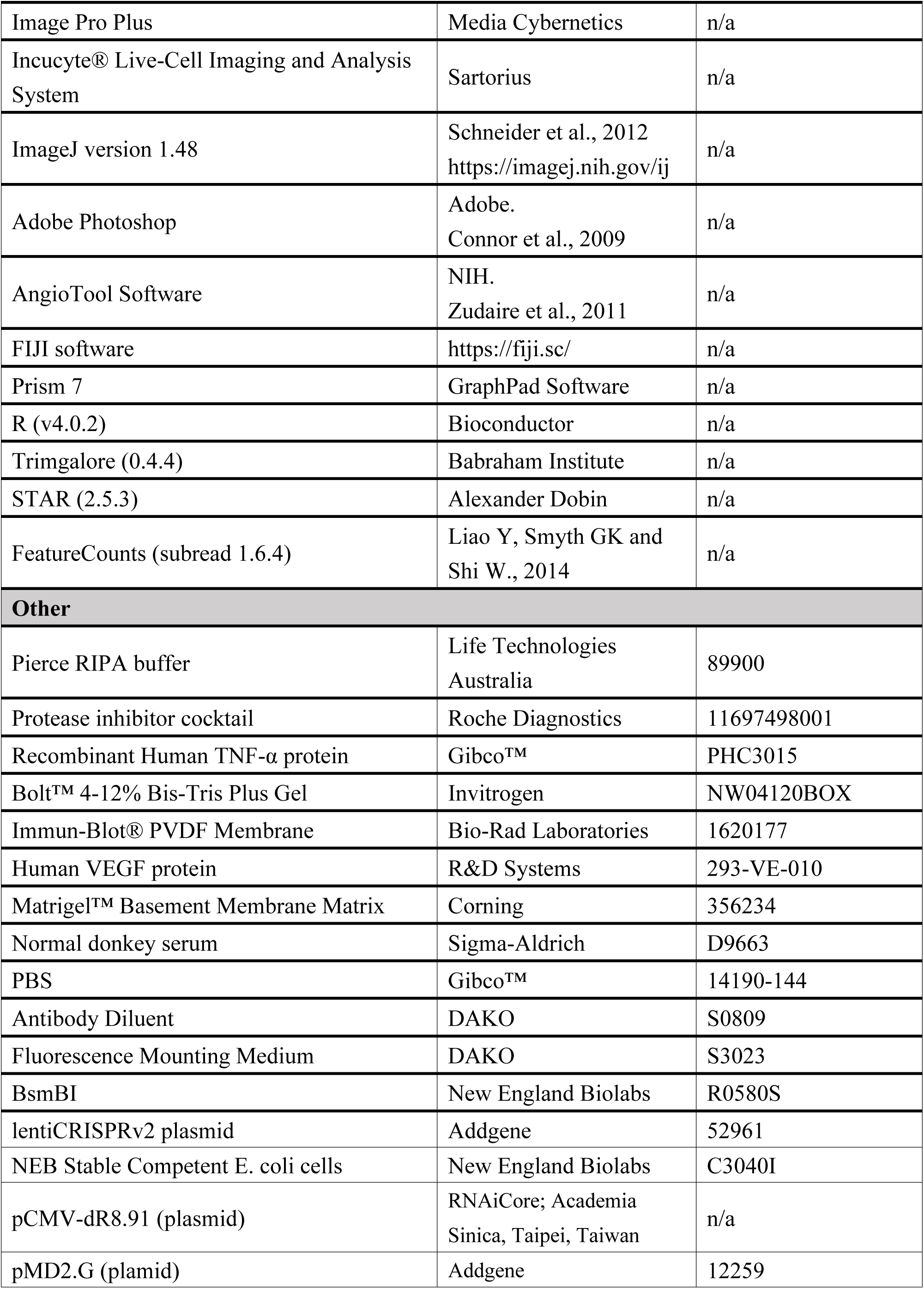
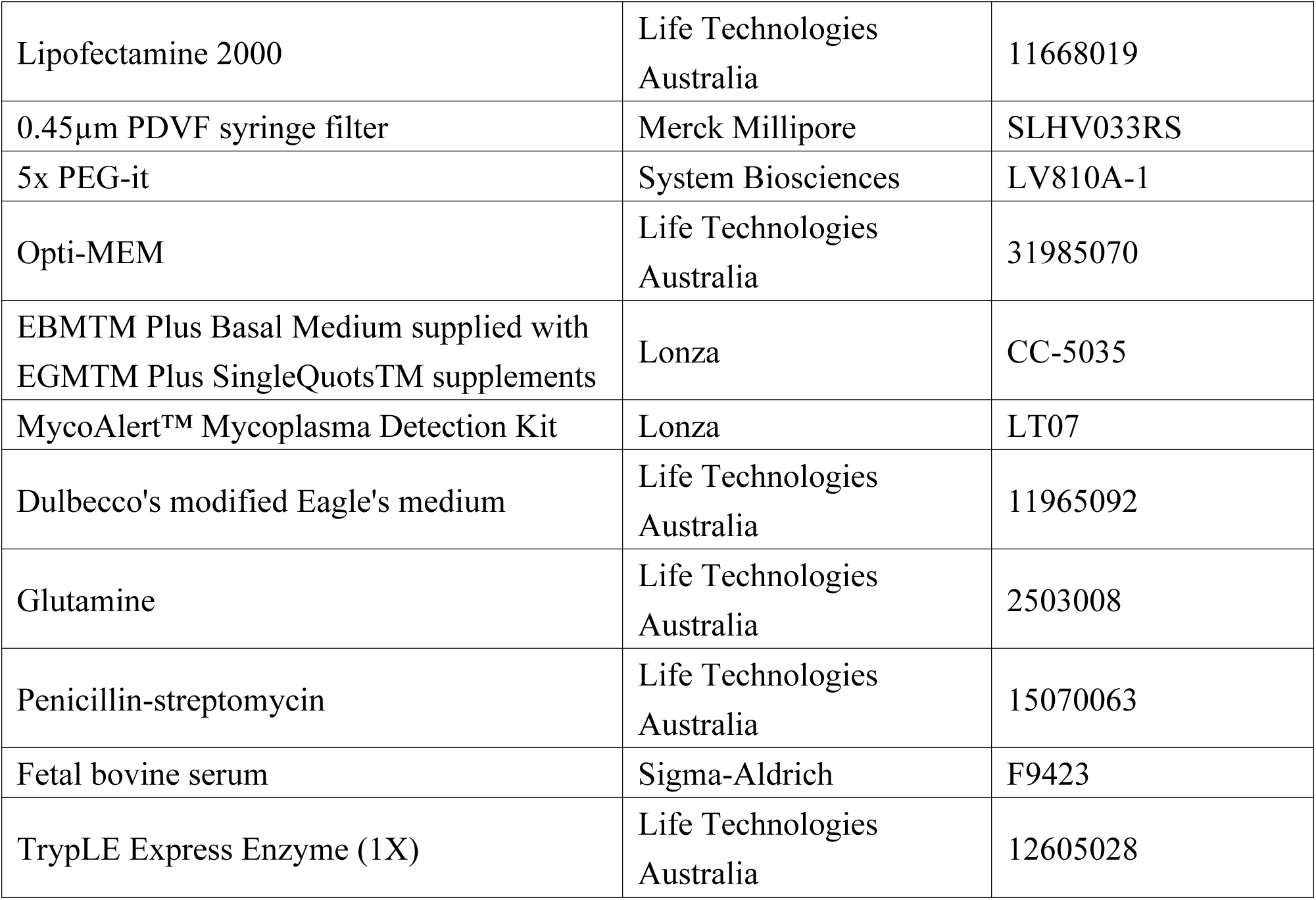
Key Resources table.

### SUPPLEMENTARY METHODS

#### Generation of TAK1-KO TIME cell line

Three sgRNAs targeting the exonic regions of the human TAK1 gene (**Table S2**) were designed on CHOPCHOP (https://chopchop.cbu.uib.no). The annealed oligos ordered from the Integrated DNA Technologies (IDT; Coralville, Iowa, USA) representing the various sgRNAs were cloned into the BsmBI sites (R0580S; New England Biolabs) of the lentiCRISPRv2 plasmid (52961; Addgene). The plasmid was validated by Sanger sequencing with 3500 Series Genetic Analyzers (Thermo Fisher Scientific; Waltham, MA). HEK293FT cells (R70007; Life Technologies) were plated at 1×10^7^ cells per T75 flask one day prior to transfection. Lentivirus was made by co-transfecting lentiCRISPRv2 (15µg), pCMV Δ8.91 (15µg) and pMDG plasmids (1.5µg) (RNAiCore; Academia Sinica, Taipei, Taiwan) in HEK293FT cells using Lipofectamine 2000 (11668019; Life Technologies). Media was changed 5 hours post transfection. The virus-containing media was harvested and filtered through a 0.45 µm PDVF syringe filter (SLHV033RS; Merck Millipore) to remove cell debris 48 hours post transfection. Lentivirus aliquots were precipitated by 5x PEG-it (LV810A-1; System Biosciences) at 4°C overnight and centrifugated at 1500g at 4°C for 30 minutes. After discarding the supernatant, the lentivirus were resuspend within opti-MEM ready for use. Wild type TIME cells (7.5×10^5^) were plated at T75 flask before transduction. When grow to 50% confluence, cells were transduced by TAK1 sgRNA lentiCRISPRv2 lentivirus with polybrene (8µg/mL) (H9268; Sigma-Aldrich) for 72 hours. Medium containing 0.5 µg/mL puromycin (P8833; Sigma-Aldrich) was then replaced for another 72 hours to select stable clones which were subsequently cultured as the stable TAK1 KO cell lines.

#### Single colony selection of TAK1-KO cell line

Once the selected TAK1 KO cell lines reached 80% confluence, cells were trypsinized from the stable cell pool using TrypLE Express Enzyme (1X) (12605028; Life Technologies) to break up any clumps into individual cells. TAK1-KO cells were then diluted to a density of 5 cells/mL and resuspended with 100 ul placed in each well in 96-well plate. After expanding for 7-14 days, colonies of single cells were subsequently cultured as the single colony selected TAK1-KO cell line in culture dishes by trypsinization.

#### Identification of TAK1 indels in TAK1-KO line

Genomic DNA was isolated from wild type and TAK1-KO TIME cell lines using Quick-DNA Microprep Kit (D3021; Zymo Research) according to the manual. Extracted genomic DNA was used as a PCR template to amplify the target of the TAK1 sgRNA using sequence specific primers (**Table S2**). Briefly, PCR reactions were made up to 25 µL comprising 12.5 µL Q5 Hot Start High-Fidelity 2X Master Mix (M0494; New England Biolabs), 1.25 µL of forward and reverse primers and the diluted gDNA, under thermocycling conditions of 98°C initial denaturation for 30 seconds, and 30 cycles of 98°C denaturation for 10 seconds, 65°C annealing for 30 seconds, and 72°C extension for 30 seconds with a 72°C final extension for 2 minutes. The PCR amplification product was validated using electrophoresis using 1% agarose gel (A6361; Sigma-Aldrich). The gDNA was then cleaned by using a Quick-DNA Microprep Kit (D3021; Zymo Research). Following the first-round of PCR amplification, a second round of sequencing PCR was performed. The second-round PCR products were subsequently purified using Agencourt CleanSeq Kit (A29151; Beckman Coulter). The PCR samples were denatured at 98°C for 5 minutes and sequenced by Sanger sequencing.

#### Electroretinography (ERG) in rat

Rats were dark-adapted overnight prior to the ERG assessment under fully dark-adapted conditions. Details for functional assessment are as previously described (58). Briefly, a pair of custom-made chloride silver active and ring-shaped reference electrodes (99.9%, A&E Metal Merchants, Sydney, NSW, AU), connected to platinum leads (F-E-30, Grass Telefactor, West Warwick, RI), was place on the central cornea and around the sclera, respectively. A stainless-steel needle electrode (F-E2-30, Grass Telefactor) was inserted subcutaneously into the tail of the animal to act as the ground. Measurements were recorded simultaneously for both eyes. ERG analysis was undertaken as previously described to return measures for three major waveforms, which are known to reflects the function of key retinal cells, specifically photoreceptors (a-wave), bipolar cells (b-wave), and ganglion cells (scotopic threshold response, STR).

#### Spectral-domain optical coherence tomography (OCT) in rat

Following ERG measurement, rat eyes were imaged using Envisu R2200 VHR spectral domain-OCT (Bioptigen, Inc., Morrisville, NC, USA). Volume scans consisting of 1000 A-scans per 100 B-scans (100 horizontal B-scans evenly spaced in the 1.4 mm vertical dimension) centered on the optic nerve head (ONH; 1.4 × 1.4 × 1.57 mm) were obtained from both eyes. Four B-scans that crossed the ONH were analyzed using FIJI software (https://fiji.sc/). In each B-scan the inner limiting membrane, retinal nerve fiber layer, inner plexiform layer and Bruch’s membrane were manually segmented by a masked observer as previously described (58). Total retinal thickness was measured from the inner limiting to Bruch’s membrane. Retinal nerve fiber layer thickness was measured from the inner limiting membrane to the inner aspect of the inner plexiform layer. Outer retinal thickness was measured from Bruch’s membrane to the outer plexiform layer.

### SUPPLEMENTARY FIGURES

**Fig. S1.**
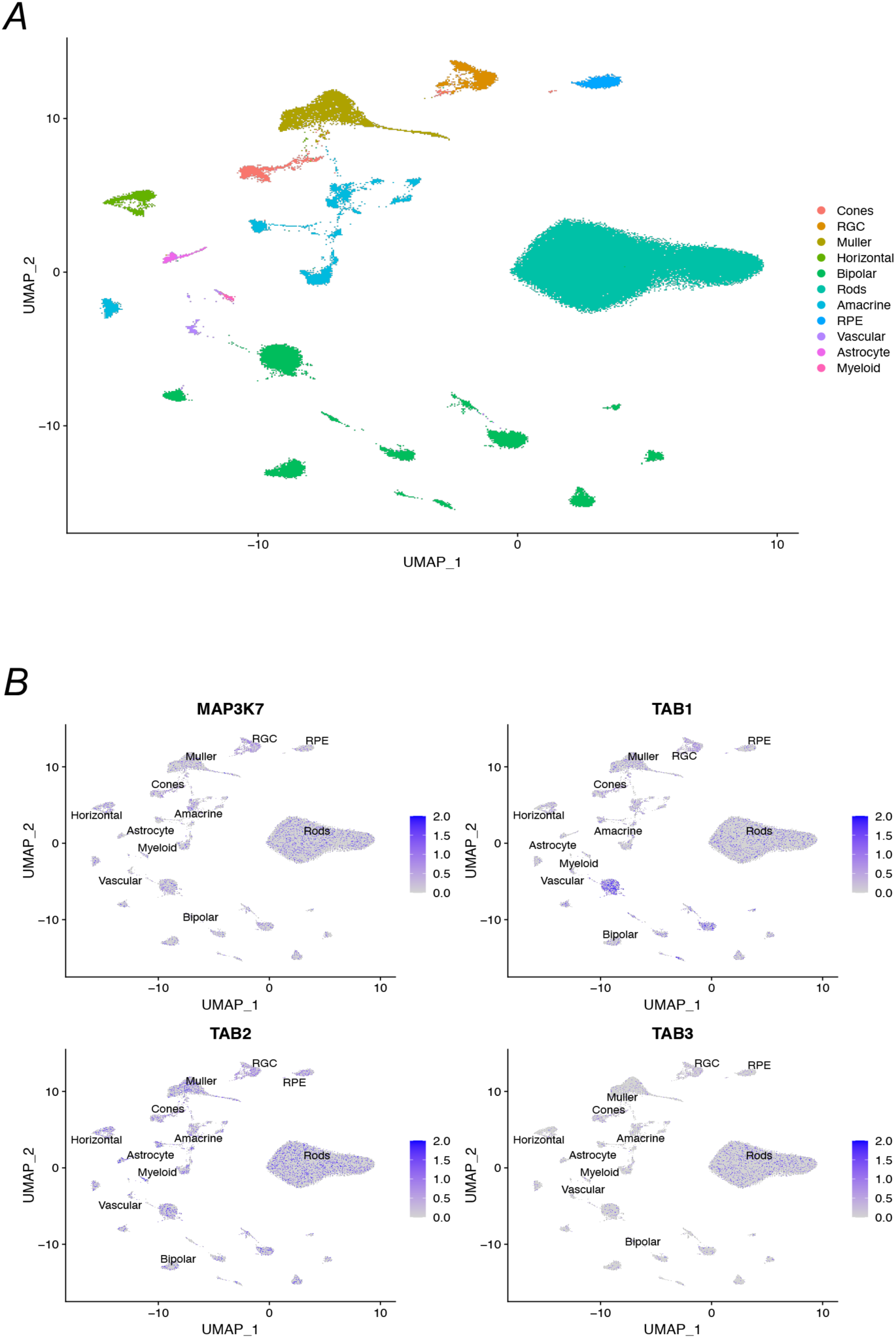
Uniform Manifold Approximation and Projection (UMAP) clustering of the integrated datasets from snRNA-seq data (A) representing the expressions of TAK1 (MAP3K7) and TAK1-associated proteins (TAB1-3) in identified retinal and non-retinal cellular clusters (B).

**Fig. S2.**
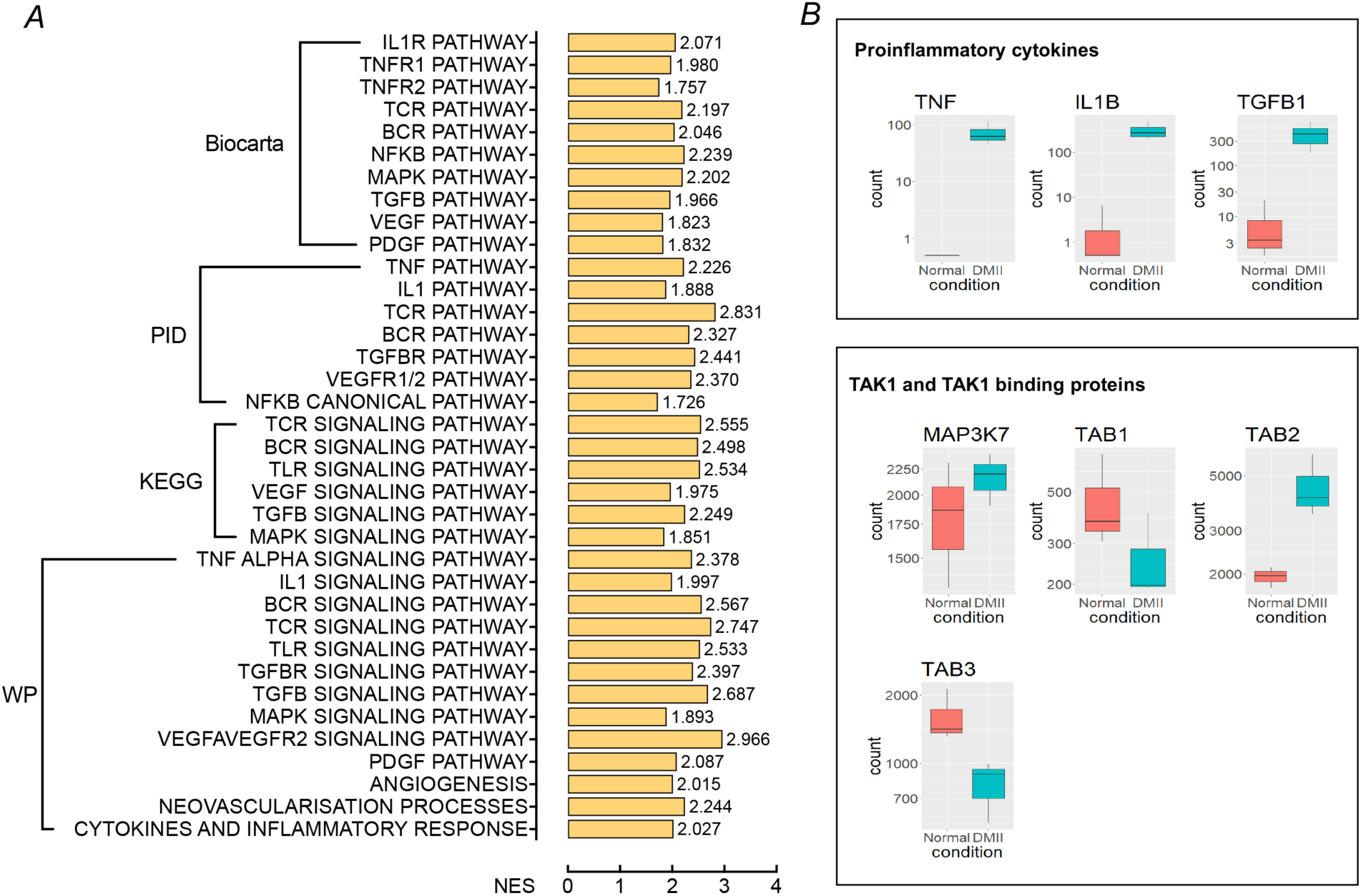
(A) Gene set enrichment analysis (GSEA) revealed positive enrichment of TAK1 upstream pathways in the retinae from patients with retinal neovascularization due to proliferative diabetic retinopathy. Gene set database includes Biocarta, Pathway Interaction Database (PID), Kyoto Encyclopedia of Genes and Genomes (KEGG) and WikiPathways (WP). (B) Readcounts of selected genes related to TAK1 in the retinae from normal control and patients with retinal neovascularization due to diabetic retinopathy. False discovery rate (FDR) was set at a q value <0.05.

**Fig. S3.**
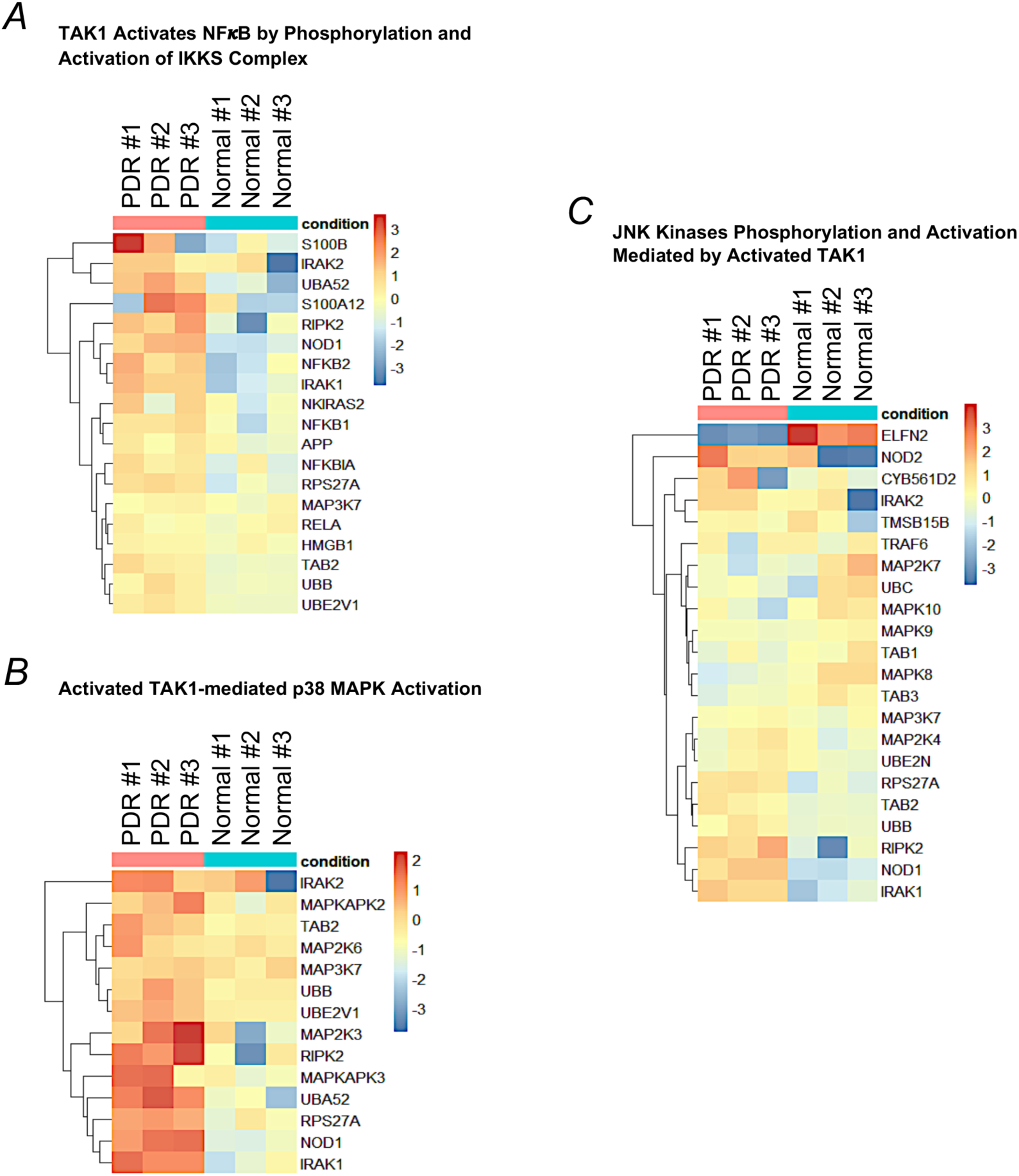
Heat maps generated from unsupervised clustering of genes from Reactome Pathway Database of (A) “TAK1 Activates NFκB by Phosphorylation and Activation of IKKS Complex”, (B) “Activated TAK1-mediated p38 MAPK Activation, and (C) “JNK kinases phosphorylation and activation mediated by activated TAK1”. False discovery rate (FDR) was set at a q value <0.05.

**Fig. S4.**
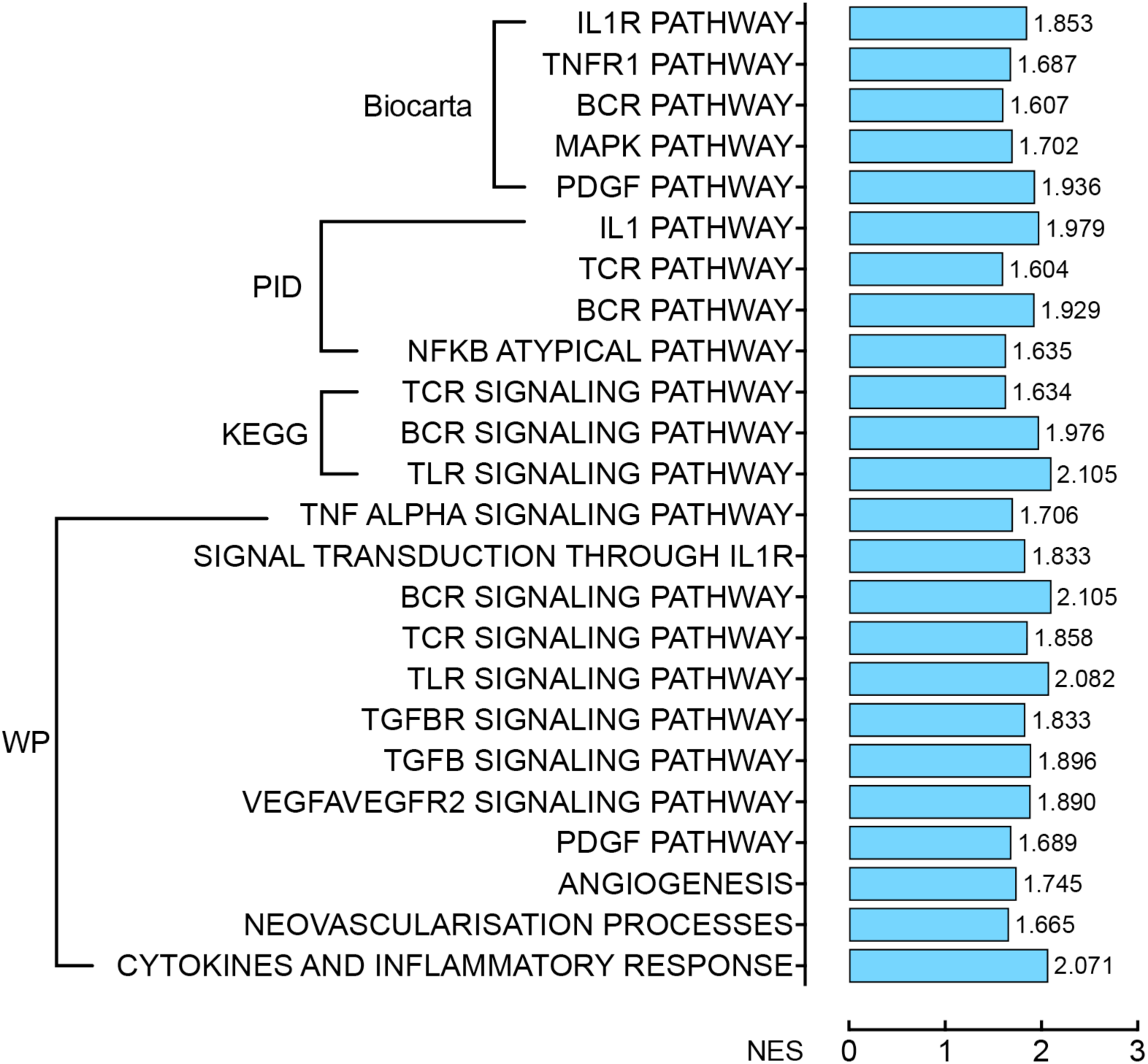
Gene-set enrichment analysis from multiple databases indicated the positive impact of TAK1 upstream pathways in rat OIR retinas. Genome-wide transcriptional analysis of rat OIR retina samples indicated the differential gene expression associated with positive impact in TAK1 upstream pathways. Biocarta, Pathway Interaction Database (PID), Kyoto Encyclopedia of Genes and Genomes (KEGG), and WikiPathways (WP) gene sets were used in GSEA. False discovery rate (FDR) was set at a q value <0.05.

**Fig. S5.**
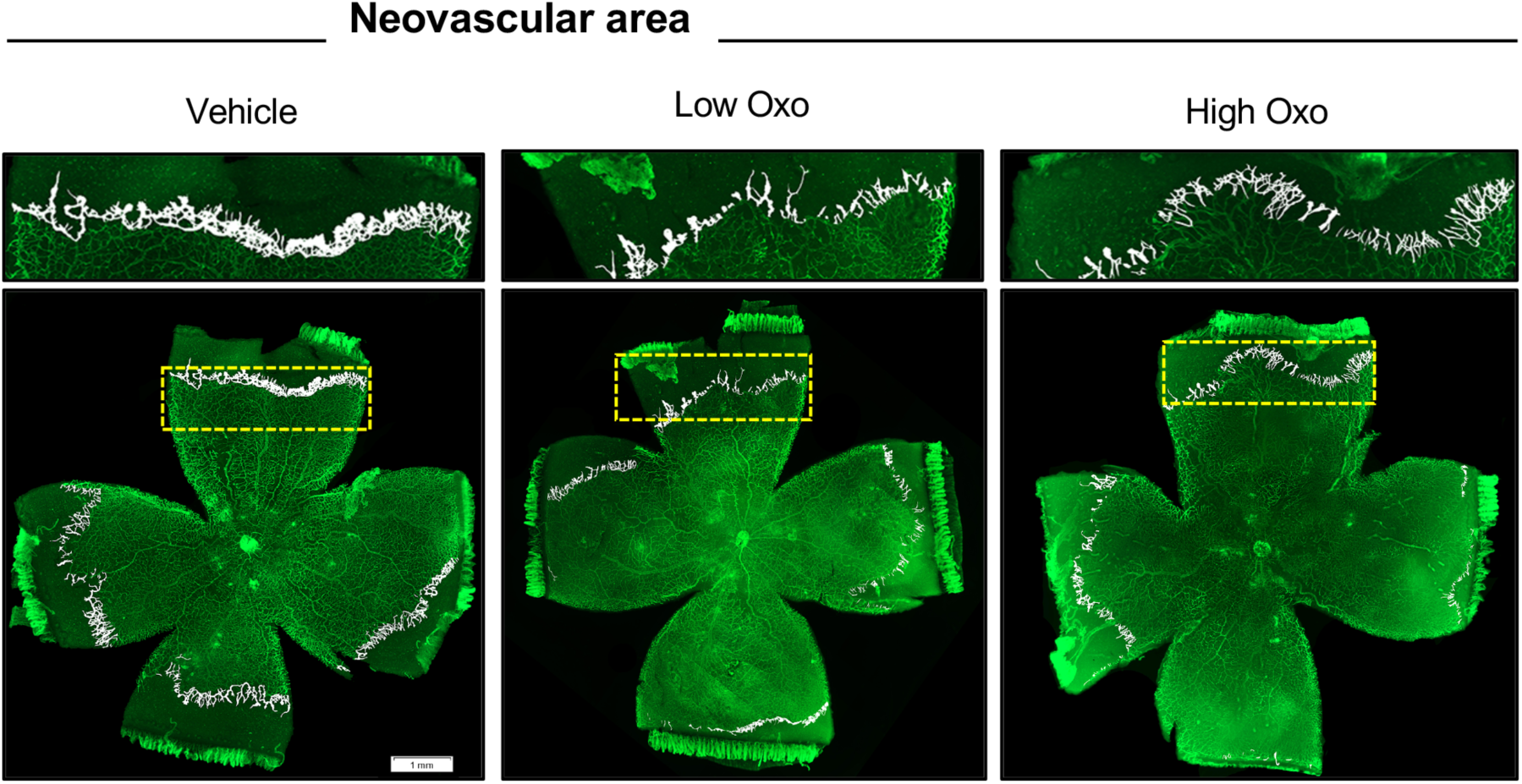
Uncropped retinal flat mounts. The region indicated by the dashed line was used to show retinal neovascularization (highlighted in white) in Fig. 3B. Oxo: 5Z-7-oxozeaenol. Scale bar: 1 mm.

**Fig. S6.**
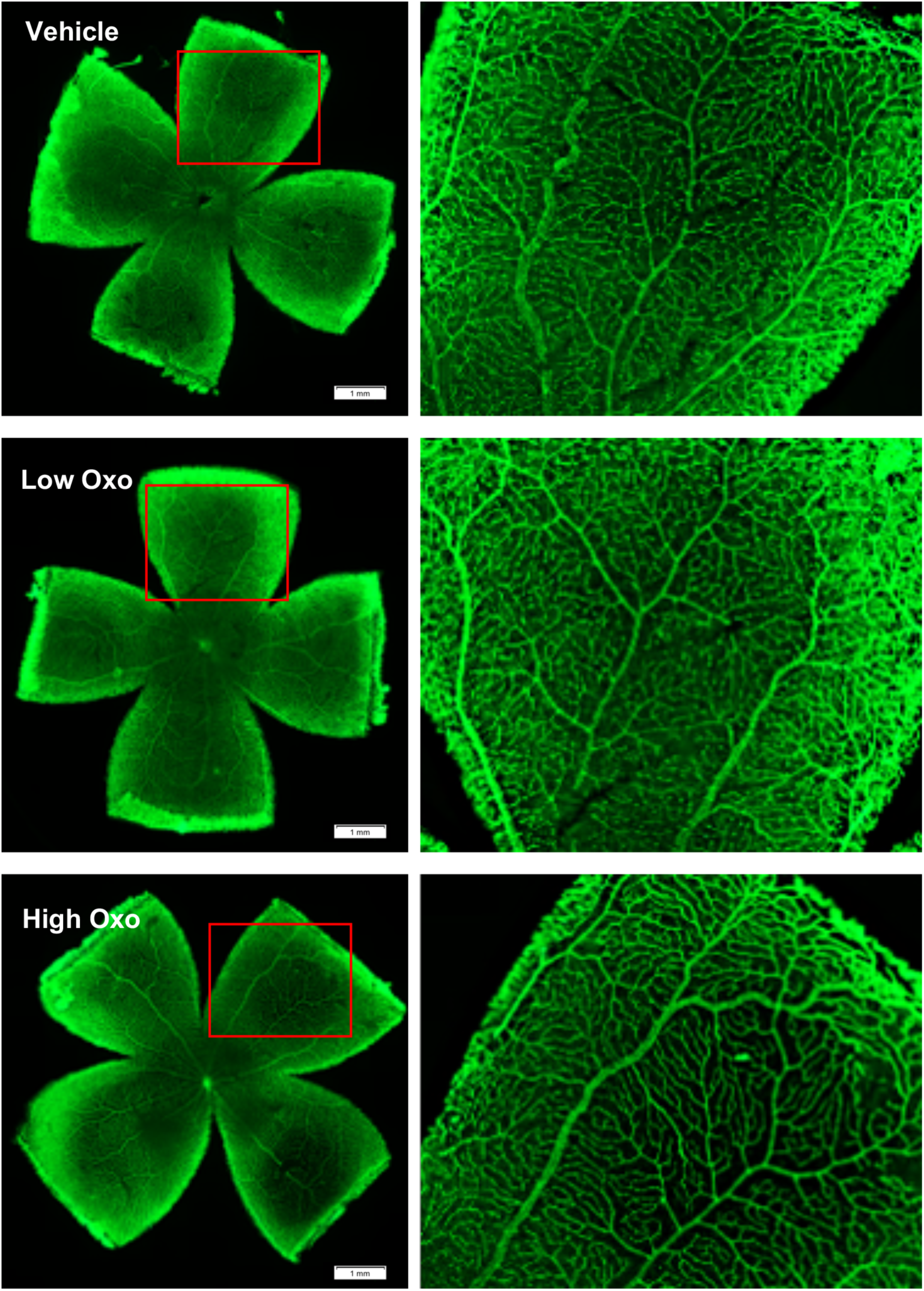
Inhibition of TAK1 did not cause retinal vessel remodeling. Retinal flat mount was labelled with IB4 28 days after a single intravitreal injection of vehicle, low (18 ng) or high (90 ng) dose of 5Z-7-oxozeaenol (Oxo) in normoxic rats to examine drug effects on normal retinal vasculature. Scale bar: 1 mm.

**Fig. S7.**
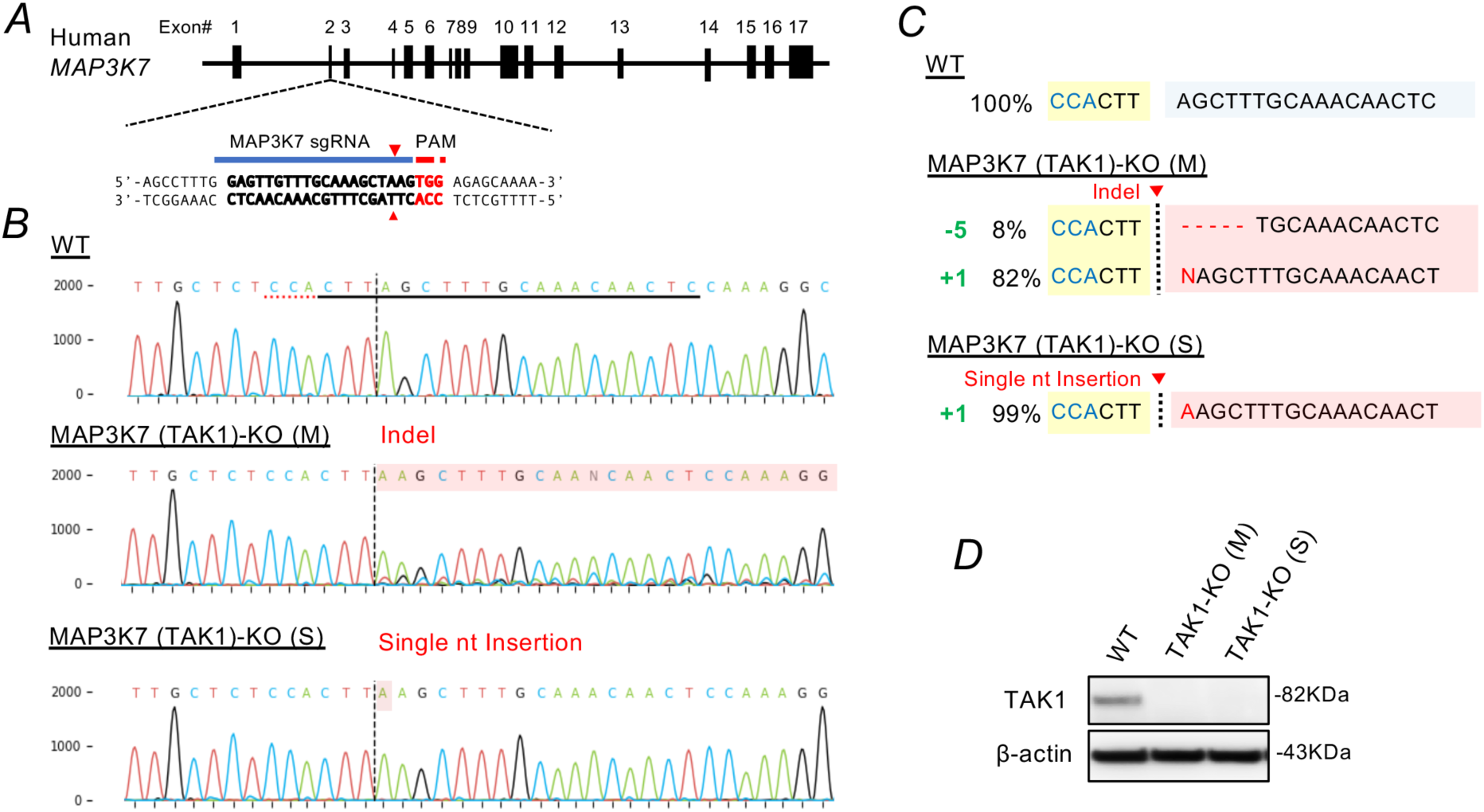
Generation of CRISPR/Cas9-mediated MAP3K7 knockout (TAK1-KO) in human endothelial cells. (A) Genome of TAK1 (MAP3K7). A sgRNA was designed to target exon 2 of the TAK1 gene. (B) As compared with wild-type (WT) TIMEs, genomic DNA sequences revealed a heterogeneous MAP3K7-KO (M) and a single homogenous MAP3K7-KO (S) TIMEs were established after CRISPR/Cas9 gene editing. (C) Sanger sequencing confirmed that MAP3K7-KO (M) TIMEs had mixed indel profiles, and MAP3K7-KO (S) TIMEs was a single nucleotide insertion, which created as premature Stop Condon in TAK1 exon 2 region. (D) Western blot characterization of TAK1 knockout in MAP3K7-KO (M) and MAP3K7-KO (S) TIMEs.

**Fig. S8.**
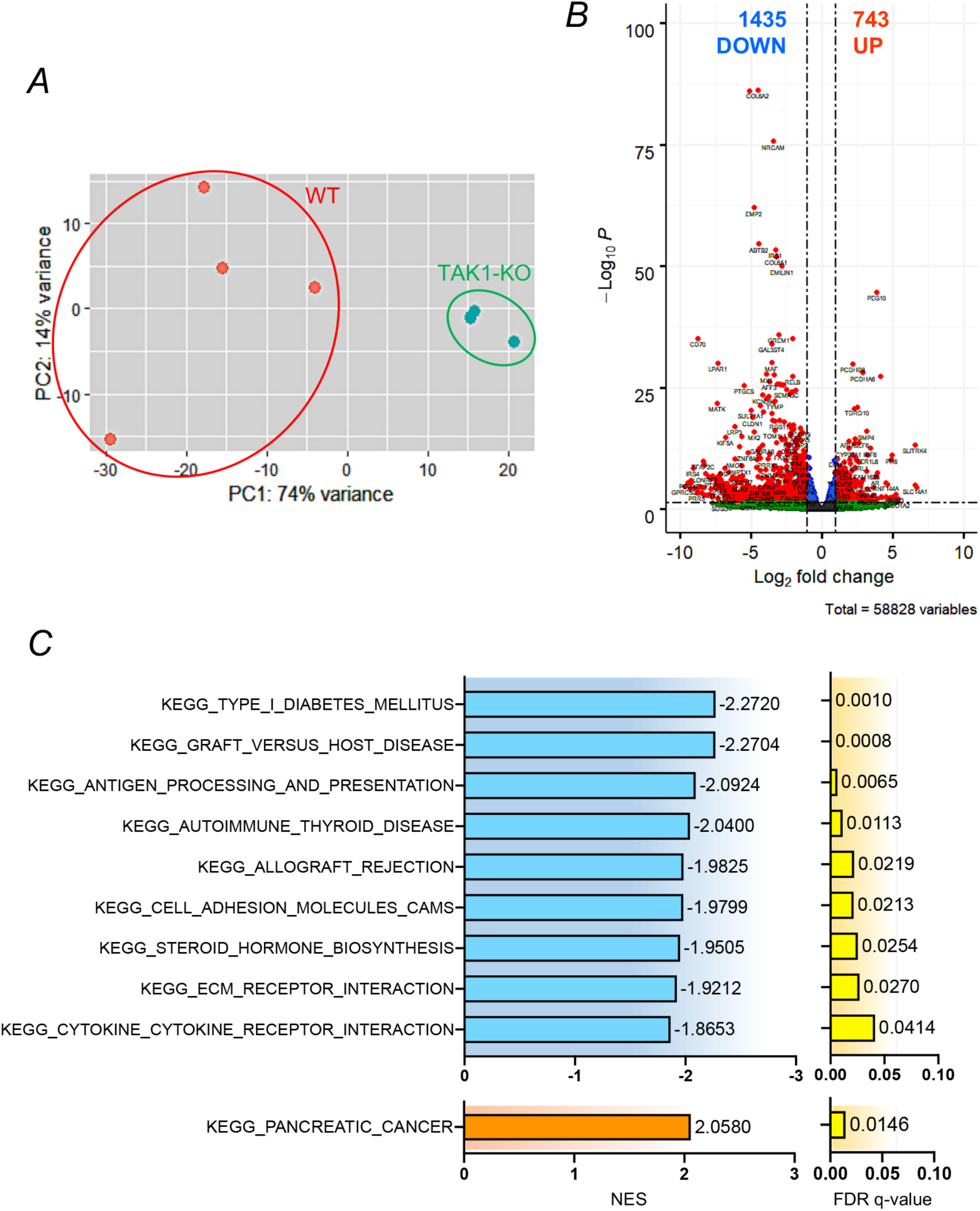
(A) Principal components analysis of RNA-seq data of wild type cells, wild-type (WT) cells and TAK1 knockout (TAK1-KO) TIMEs. Each dot represents an experimental replicate. (A) Volcano plots showing genes that were significantly different between WT and TAK1-KO TIMEs. Red dots refer to genes showing a log2|fold change| > 1 and FDR < 0.05. Labels indicate the significantly up-regulated genes in each data set. (C) Gene set enrichment analysis (GSEA) revealed the top 10 Kyoto Encyclopedia of Genes and Genomes (KEGG) pathways that were positively or negatively enriched in TAK1-KO TIMEs. False discovery rate (FDR) was set at a q value <0.05.

**Fig. S9.**
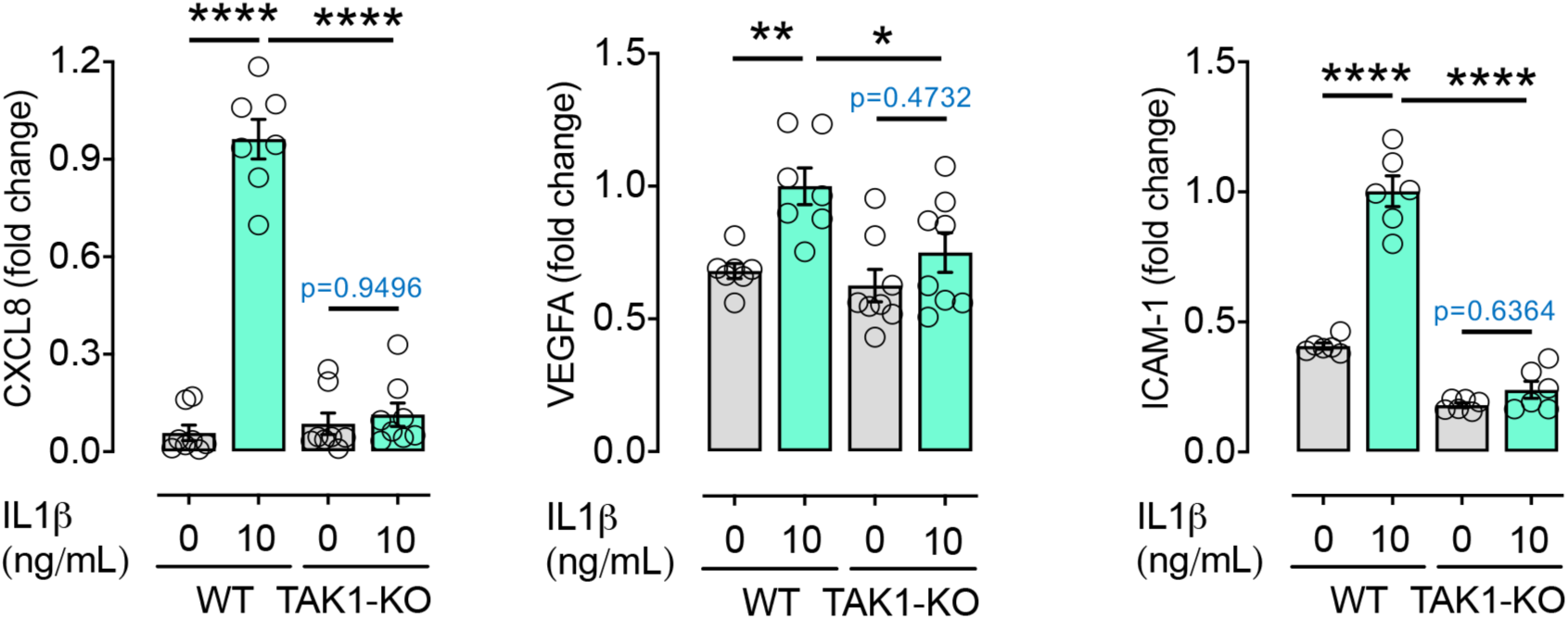
TAK1 deletion reduced the expression of *TAK1, ICAM-1, CXCL8 and VEGF* genes in human microvascular endothelial cells. qPCR results revealed that TAK1 deletion suppressed the expression of selected cytokines (*CXCL8* and *VEGFA)* and adhesion molecules (*ICAM-1)* in TIMEs stimulated with IL1*β* for 24 hours (n = 4; technical duplicate from each biological repeat was shown as separated dot plot). Group data are shown as means ± SEM. Statistical analysis was undertaken with one-way ANOVA and Tukey’s multiple comparison test; \**P* < 0.05, \*\**P* < 0.01, \*\*\**P* < 0.001, \*\*\*\**P* < 0.0001.

**Fig. S10.**
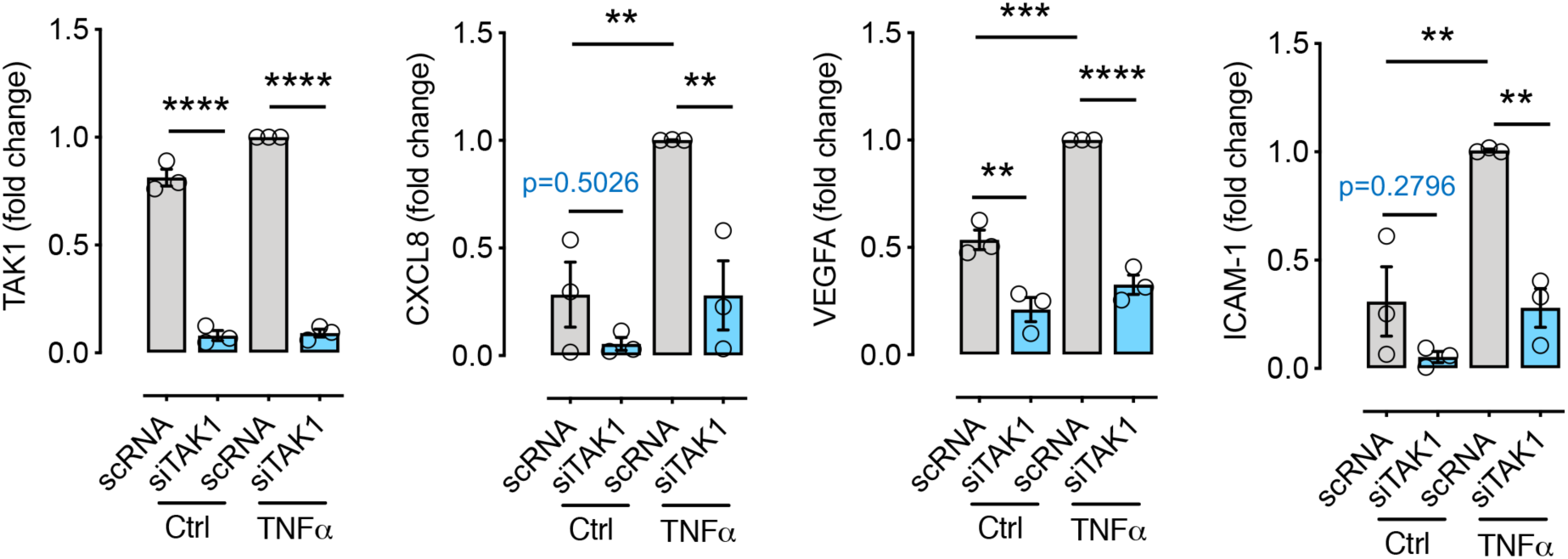
TAK1 knockdown by siRNA inhibited the expression of *TAK1, ICAM-1, CXCL8 and VEGF* genes in human primary retinal microvascular endothelial cells (HRMECs). qPCR results revealed that TAK1 knockdown by siRNA targeting TAK1 (siTAK1) suppressed the expression of selected cytokines (*CXCL8* and *VEGFA)* and adhesion molecules (*ICAM-1)* in HRMECs stimulated with TNFα for 24 hours (n = 3). Group data are shown as means ± SEM. Statistical analysis was undertaken with one-way ANOVA and Tukey’s multiple comparison test; \*\**P* < 0.01, \*\*\**P* < 0.001, \*\*\*\**P* < 0.0001.

**Fig. S11.**
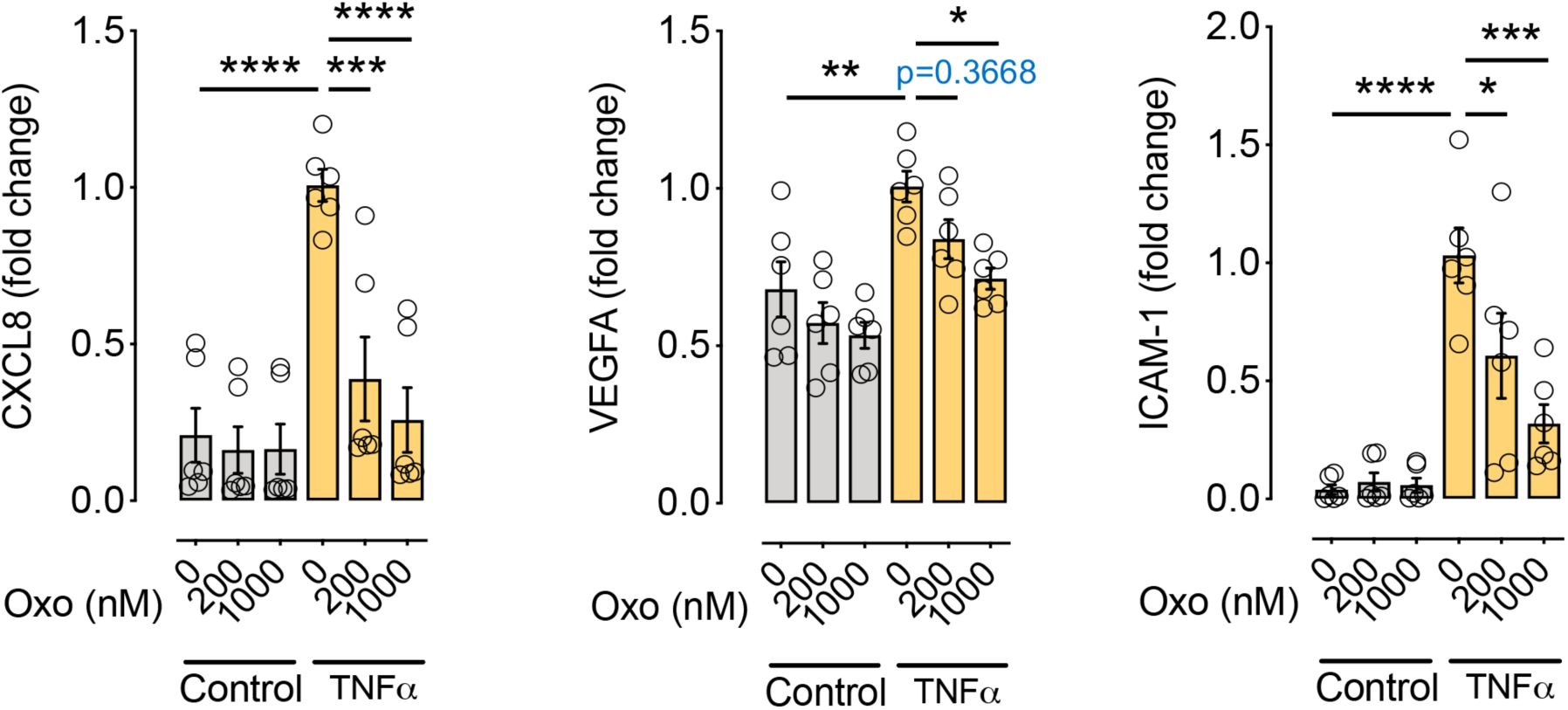
TAK1 inhibition by 5Z-7-oxozeaenol inhibited the expression of *TAK1, ICAM-1, CXCL8 and VEGF* genes in primary human retinal microvascular endothelial cells (HRMECs). qPCR results revealed that pharmacological inhibition of TAK1 by 5Z-7-oxozeaenol (Oxo) suppressed *ICAM-1*, *PTGS2*, *CXCL8* and *VEGFA* in HRMECs stimulated with TNFα in a dose-dependent manner (n = 3; technical duplicate from each biological repeat was shown as separated dot plot). Group data are shown as means ± SEM. Statistical analysis was undertaken with one-way ANOVA and Tukey’s multiple comparison test; \**P* < 0.05, \*\**P* < 0.01, \*\*\**P* < 0.001, \*\*\*\**P* < 0.0001.

**Fig. S12.**
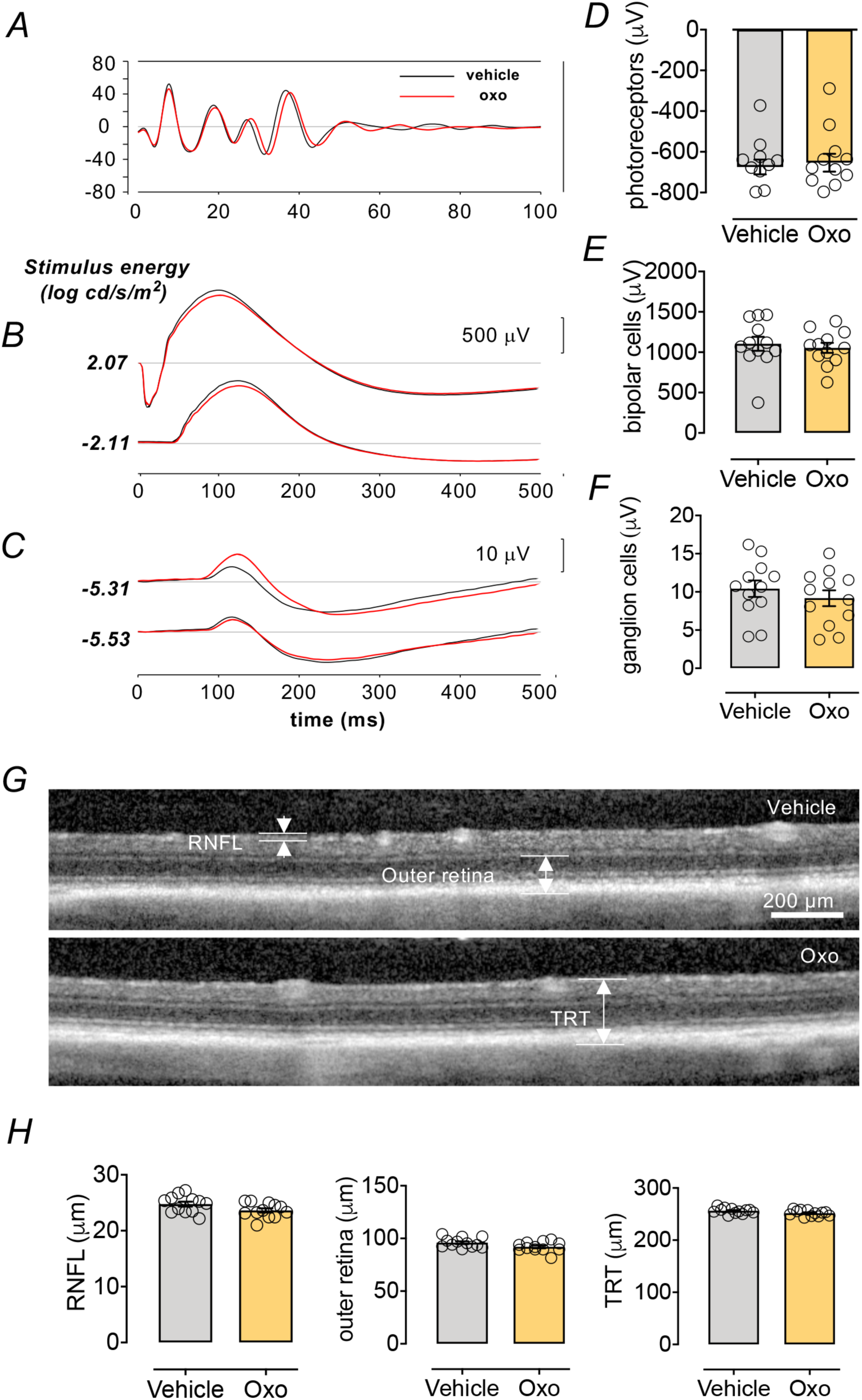
Effect of 5Z-7-oxozeaenol (Oxo) on *in vivo* retinal structure and function. ERG and OCT were performed 28 days after intravitreal injection of 5Z-7-oxozeaenol (90 ng) in normoxia Brown Norway rats. No difference was observed between the vehicle- (black traces) and 5Z-7-oxozeaenol-injected (red traces) eyes in terms of the average amacrine cell oscillatory potentials (A) photoreceptoral and bipolar cell responses (B) or ganglion cell responses (C) in 5Z-7-oxozeaenol-injected eyes (n = 11, red traces) and corresponding vehicle-injected eyes (n = 12, black traces). Group average (± SEM) photoreceptor (D. *P* = 0.71) bipolar cell (E. *P* = 0.63) and ganglion cell (F, *P* = 0.41). Response amplitude were not different between 5Z-7-oxozeaenol-injected eyes and corresponding vehicle-injected eyes. (G) Representative OCT images from a 5Z-7-oxozeaenol- and vehicle-injected eye. (H) Total retinal thickness (TRT), outer retina, and retinal nerve fiber layer (RNFL) thickness were not different between groups (*P* = 0.0639 of TRT, *P*=0.0502 of outer retina, *P* = 0.064 of RNFL). All data are shown expressed as means ± SEM. Statistical analysis was undertaken with two-tailed Student’s *t*-test.

